# Engineering a Scalable and Orthogonal Platform for Synthetic Communication in Mammalian Cells

**DOI:** 10.1101/2023.01.18.524631

**Authors:** Anna-Maria Makri Pistikou, Glenn A.O. Cremers, Bryan L. Nathalia, Bas W.A. Bögels, Bruno V. Eijkens, Anne de Dreu, Maarten T.H. Bezembinder, Oscar M.J.A. Stassen, Carlijn C.V. Bouten, Maarten Merkx, Roman Jerala, Tom F. A. de Greef

**Affiliations:** Laboratory of Chemical Biology, Department of Biomedical Engineering, Eindhoven University of Technology, Eindhoven, The Netherlands; Institute for Complex Molecular Systems, Department of Biomedical Engineering, Eindhoven University of Technology, Eindhoven, The Netherlands; Computational Biology Group, Department of Biomedical Engineering, Eindhoven University of Technology, Eindhoven, The Netherlands; Laboratory for Cell and Tissue Engineering, Department of Biomedical Engineering, Eindhoven University of Technology, Eindhoven, 5612 AZ, the Netherlands; Department of Synthetic Biology and Immunology, National Institute of Chemistry, Ljubljana, Slovenia; EN-FIST Centre of Excellence, Ljubljana, Slovenia; Institute for Molecules and Materials, Radboud University, Nijmegen, The Netherlands; Center for Living Technologies, Eindhoven-Wageningen-Utrecht Alliance, the Netherlands

## Abstract

The rational design and implementation of synthetic, orthogonal mammalian communication systems has the potential to unravel fundamental design principles of mammalian cell communication circuits and offer a framework for engineering of designer cell consortia with potential applications in cell therapeutics and artificial tissue engineering. We lay here the foundations for the engineering of an orthogonal, and scalable mammalian synthetic intercellular communication platform that exploits the programmability of synthetic receptors and selective affinity and tunability of diffusing coiled-coil (CC) peptide heterodimers. Leveraging the ability of CCs to exclusively bind to a selected cognate receptor, we demonstrate orthogonal receptor activation, as well as Boolean logic computations. Next, we reveal synthetic intercellular communication based on synthetic receptors and secreted multidomain CC ligands and demonstrate a minimal, three-cell population system that can perform distributed AND gate logic. Our work provides a modular and scalable framework for the engineering of complex cell consortia, with the potential to expand the aptitude of cell therapeutics and diagnostics.

## Introduction

The ability to engineer novel functions in mammalian cells has become a key force in biomedical research, revolutionizing the field of cell-based diagnostics and therapeutics^1–4^. In particular, the introduction of synthetic receptors^5,6^ has enabled the engineering of designer cells that, due to their sensing and actuating capabilities, can detect and correct disease state^7–16^. To further improve on the specificity of such approaches, multi-layered circuits for combinatorial detection of multiple biomarkers need to be introduced^17–19^. Nonetheless, large genetic circuits are challenging to implement in monocultures of cells, as they are met with limitations due to the burden introduced by resource sharing^20–22^. Additionally, delivery of multiple genetic constructs can be further restricted due to the limited effective packaging capacity of the vector^23^. To fully unlock the potential of such a technology, engineered cells need to communicate reciprocally and aptly process information in a way that resembles specialised cell consortia of the human immune system. The human immune system is a compelling model target, since it has the exceptional capacity to sense and respond to a multitude of diffusing signals, perform distributed computing, and leverage specialised cell types to address challenges. An envisioned mammalian synthetic communication network has the potential to reduce burden due to resource competition on individual cell populations by permitting distributed information processing, leading to advanced functions^24,25^. Such a system has the capacity to enable population control as well as allow for coordinated responses of therapeutic cells, in a manner similar to previously developed synthetic prokaryotic communication systems^26–28^. To realize intercellular communication and enable programmable control of neighbouring cell responses, ideally cells are equipped with the capacity to sense orthogonal, soluble stimuli while at the same time producing user-defined responses. Synthetic, intercellular, juxtracrine communication between mammalian cells has been previously achieved utilizing synthetic receptors^12,15,29^ that can perform logic operations^17– 19,30,31^ based on cell-to-cell contact. Efforts to introduce diffusion-based, intercellular synthetic communication in mammalian cells include the design of mammalian circuitry deploying repurposed small molecules^32–37^, in addition to approaches that employ directed evolution of naturally occurring polypeptides^38^. Although such approaches have managed to render cells capable of complex biocomputations^39^, they lack in scalability due to the use of small molecules as signal initiators or require labour intensive methods to render the input signal orthogonal. While these advancements have arguably managed to move mammalian synthetic biology forward, there is currently no available platform for synthetic communication between mammalian cells, utilising diffusible ligands, that is inherently scalable, and orthogonal to the native cell.

Here, we engineer a scalable and orthogonal synthetic communication platform for mammalian cells. In detail, we functionalised erythropoietin receptor (EpoR) domains from the Generalized Extracellular Molecule Sensor (GEMS) platform^10^ with coiled-coil peptides (CC) from the NICP set^40,41^, that possess the ability to mutually and orthogonally bind to each other (Figure 1a). Programmable design of large sets of CC heterodimers has been achieved^42^, providing our platform with potential scalability. The similarity in peptide size and conformation amongst individual CC interaction domains generalizes the receptor activation mechanism and hence allows for the introduction of different CC modules on the EpoRs without having to re-engineer the system for each orthogonal pair of the ligand-receptor separately. We first demonstrate that receptors carrying heterodimeric, complementary CC peptides result in robust receptor activation upon expression. The design and engineering of ditopic CC peptide ligands allows for cognate receptor activation by addition of the ditopic ligands to cell culture. The present work exemplifies the scalability of our platform by showcasing synthetic receptor activation by three unique CC pairs. Additionally, we demonstrate that mammalian cells expressing a combination of synthetic receptors can perform two-input logic (AND and OR gate operations) (see Figure 1b). Finally, we achieve synthetic intercellular communication in a minimal system composed of a sender population that secretes a synthetic dipeptide ligand and a receiver population expressing the cognate receptor and reporter. Establishing an inducible intercellular communication system ultimately allows for the engineering of a three cell-population system that can perform distributed logic operations. Our design offers a customizable and extensible framework facilitating the engineering of intricate consortia of designer cells that can process information and autonomously respond to stimuli. We propose that the present work holds the potential to expand the aptitude of cell therapeutics and diagnostics, but also offers a platform to study fundamental principles of mammalian cell-cell communication circuits.

**Fig. 1:**
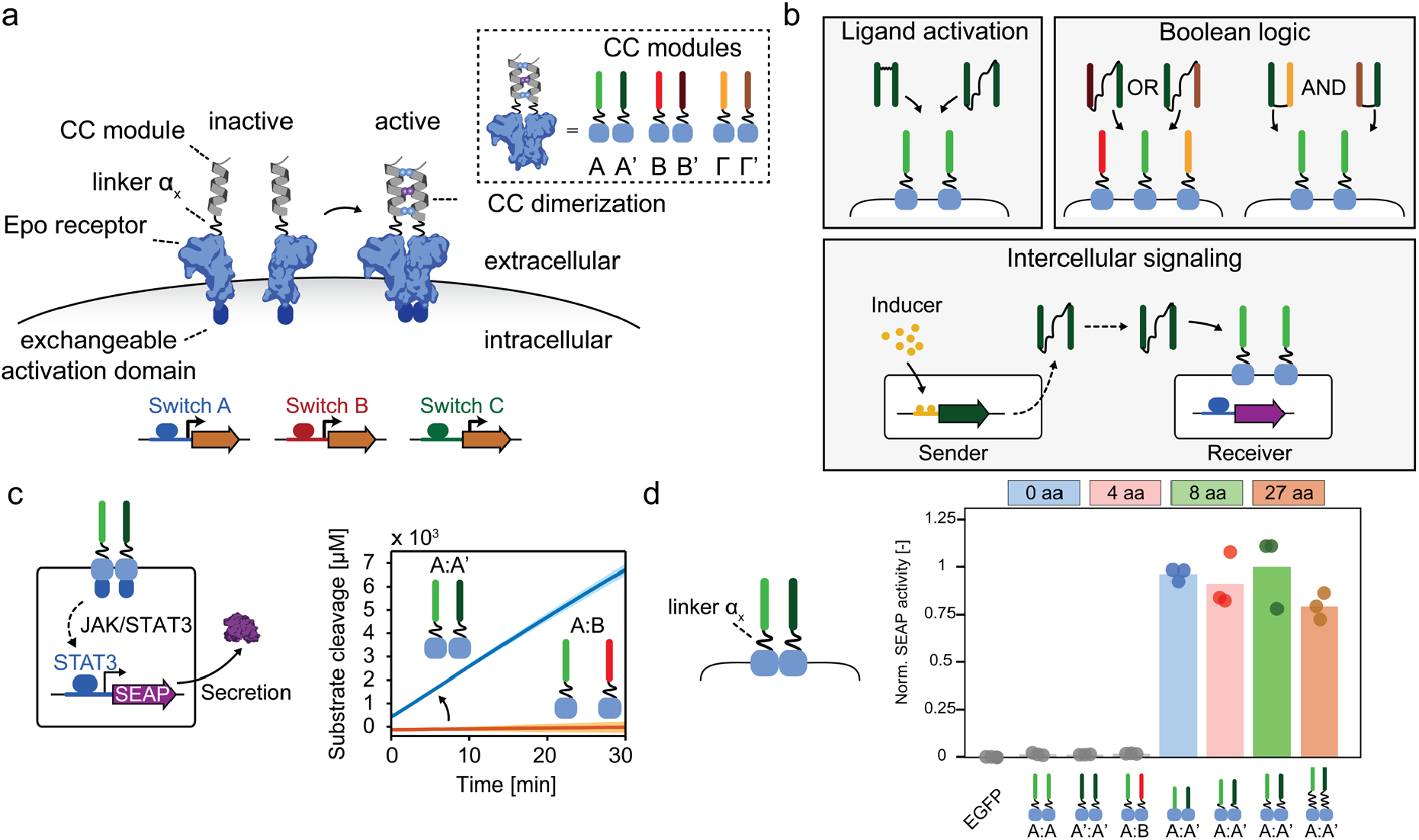
Design elements and general overview of a coiled-coil (CC) functionalised-GEMS synthetic communication platform for mammalian cells. **a** Schematic representation of CC-functionalised GEMS receptor resulting in activation of target genes, upon dimerization. CC peptides are N-terminally fused, through linker **α**_**x**_ to the extracellular and transmembrane domains of the erythropoietin receptor (EpoR) cluster that can induce transgene expression following activation of an intracellular signalling pathway. Complementary CC modules are indicated as A:A’, B:B’ and Γ:Γ’ (see Supplementary Table S1). Downstream receptor signalling can be modulated by exchanging the intracellular activation domains (indicated here as switch A, B and C) and desired output can be expressed by replacing the reporter gene to a transgene of choice. **b** Receptor activation can be achieved by soluble ditopic CC ligands (upper-left panel) the properties of which can be utilised to achieve Boolean logic gate operators (AND/OR – upper-right panel). Intercellular communication (bottom panel) can be achieved by engineering sender cells that express the ligand under the control of an ON/OFF switch and a receiver cell population expressing the cognate receptor and reporter genes. **c** Schematic overview of the reporter system to monitor receptor activation using JAK/STAT (Janus kinase/signal transducer and activator of transcription) intracellular signalling (left panel). Each receptor monomer bears a cognate CC (A and A’ with linker **α**_**2**_) that can cause the receptor to heterodimerize. Phosphorylated STAT3 results in the production of the reporter protein: human placental secreted alkaline phosphatase (SEAP). SEAP catalyses the hydrolysis of a chromogenic substrate (p-Nitrophenylphosphate (pNPP) (see Methods and Supplementary Figure S1). Substrate conversion is only observed when the cognate receptors A:A’ are expressed on the cell surface, while non-cognate pairs (A:B) do not result in receptor activation and subsequent substrate conversion (right panel). The experiment is performed in independent triplicates; solid line indicates the mean and shaded area indicates standard deviation. **d** Normalised SEAP activity in HEK293T cells transiently transfected with receptor pairs with varied linker lengths **α**_**x**_ of zero, four, eight and 27 amino acids (aa; GS or GSS repeats, Supplementary Table S2 and Supplementary Figures S2 and S3b). SEAP activity was measured 48 hours following transfection (see Methods). Only the correct combination of cognate CCs (A:A’) results in receptor activation, while non-cognate pairs (A:A, A’:A’ and A:B) result in no activation. Bars indicate mean activity; individual data points represent independent triplicates. SEAP activity is normalised with respect to the SEAP activity of the A:A’ receptor heterodimer with linker **α**_**2**_.

## Results and Discussion

### Design principles of a coiled-coil functionalised-GEMS synthetic communication platform

To actualize synthetic communication in mammalian cells, we focused our attention on the previously established Generalized Extracellular Molecule Sensor (GEMS) platform^10^, that allows for customised input and output. GEMS enables the engineering of modular receptor designs that integrate user-defined, soluble ligand sensing through the introduction of a variety of engineered ligand-binding extracellular domains^10,43–45^ to transgene expression via the rewiring of distinct endogenous signalling pathways. In detail, GEMS receptors consist of a standard transmembrane scaffold, erythropoietin receptor (EpoR) that is engineered to be inert to erythropoietin, fused to intracellular signalling domains, derived from the cytokine receptor chain interleukin 6 receptor B (IL-6RB), the fibroblast growth factor receptor 1 (FGFR1), or the vascular endothelial growth factor receptor 2 (VEGFR2). GEMS dimerization leads to downstream signalling by activating the Janus kinase/signal transducer and activator of transcription (JAK/STAT) pathway (induced by IL-6RB), the mitogen-activated protein kinase (MAPK) pathway (induced by FGFR1) or the phosphatidylinositol 3-kinase/protein kinase B (PI3K/Akt) pathway (induced by VEGFR2). Transgene expression is achieved by rerouting the abovementioned pathways using minimal promoters engineered to be responsive to specific pathways^10^. We functionalised the extracellular part of the EpoR transmembrane scaffold of the GEMS receptor with coiled-coil peptides (CC), that possess the ability to mutually and orthogonally bind to each other^40,41^ (Figure 1a). Receptor heterodimerization is thus induced by cognate CC pairing; A binds exclusively to A’, B to B’ and Γ to Γ’ (see Supplementary Table S1). Each CC is genetically fused to an EpoR domain through linker **α**_**x**_ (see Supplementary table S2 and Methods). Non-cognate CC-GEMS receptors should not activate intracellular signalling, while cognate CC pairing is expected to result in transgene expression due to receptor heterodimerization.

To monitor CC induced receptor activation, HEK293T cells were transiently transfected with CC-GEMS receptors utilizing the JAK-STAT pathway (Supplementary Figure S1), as well as STAT3, and the reporter gene STAT3-induced secreted alkaline phosphatase (SEAP; see Methods and Supplementary Table S3). SEAP secretion was quantified in the cell supernatant using a colorimetric assay that reports substrate conversion (see Methods, Figure 1c and Supplementary Figure S2). When cells were co-transfected with A-type and A’-type cognate receptors with linker **α**_**2**_ of 8 aa (4x glycine-serine (GS) repeats), a sharp increase in substrate conversion was observed, indicating receptor activation. As hypothesised, non-cognate receptor pairs (A:B) did not result in receptor activation (see Figure 1c). Next, we investigated whether receptor activation is dependent on the linker length between the CC domain and the transmembrane scaffold. We designed constructs with varying linker lengths **α**_**x**_ (see Supplementary Table S2) of zero, four (**α**_**1**_; (GS)_2_), eight (**α**_**2**_; (GS)_4_) and 27 aa (**α**_**3**_; (GSS)_9_), tethering the CC and EpoR domains. Subsequently, HEK293T cells were transfected with a range of CC-GEMS pairs, STAT3 and reporter gene and SEAP activity was quantified in the cell supernatant (see Methods). To determine background alkaline phosphatase activity and assess transfection efficiency, HEK293T cells were transiently transfected with EGFP. The transfection efficiency was estimated to be 62.1% (see Supplementary Figure S3a) and the background alkaline phosphatase activity was negligible (see Figure 1d and Supplementary Figure S3b). Our data reveals that receptor activation is not critically dependent on linker length between the CC and EpoR domain of the receptor, as all cognate heterodimer receptor pairs (A:A’) resulted in robust SEAP expression (Figure 1d and Supplementary Figure S3b), with no apparent linker-dependent effect. All non-cognate receptor pairs (A:A, A’:A’ and A:B with linker **α**_**2**_) showed negligible receptor activation, verifying the orthogonality of CC pairs, and thus demonstrating the programmability of our approach. Collectively, these results indicate that CC interactions can be used as a robust, programmable tool for synthetic receptor activation.

### Design of soluble ditopic CC ligands to activate CC-GEMS receptors

Having established that cognate CC pairs are able to induce receptor hetero-dimerization, we next aimed to assess if CC-GEMS receptor dimerization and activation can be achieved by soluble, synthetic, CC ligand dipeptides. Taking into account the parallel orientation of the bound CC cognate pairs^40^, we engineered A’-A’ dipeptides that are N-termini linked (Figure 2a). To that end, we modified the A’ peptide sequence to include a single cysteine at the N-terminus (see Figure 2b, Supplementary Figure S4a and S4b). As recombinant expression of short peptides in *E. coli* can result in protein degradation or the formation of inclusion bodies^46,47^, we increased the expression and solubility of the peptide by fusion to a small ubiquitin-like modifier (SUMO) tag^48,49^. We expressed SUMO-A’ fusion protein in *E. coli* (see Methods and Supplementary Figure S5a). Successful expression and purification of the recombinant protein was confirmed, using sodium dodecyl sulfate-polyacrylamide gel electrophoresis (SDS-PAGE; Methods and Supplementary Figure S4c) and subsequent SUMO cleavage and purification resulted in the recovery of the cysteine-functionalised A’ peptide (Supplementary Figure S4d). We employed the sulfhydryl moiety of the cysteines to synthesize N-termini linked ditopic A’-A’ ligand, using a bifunctional maleimide linker (see Methods and Figure 2c top). SDS-PAGE gel analysis illustrated the successful formation of A’-A’ dipeptides and subsequent purification with anion exchange chromatography resulted in the removal of unreacted A’ monomers (Figure 2c bottom). To validate that dipeptide formation occurs due to reaction of sulfhydryl moieties of A’ monomers, we incubated A’ monomers that were site-specifically mutated to include a glycine in place of cysteine (A’.C4G (**v**_**1**_); Supplementary Table S4, Supplementary Figure S6a and S6b) with the bifunctional linker and observed no dipeptide formation (Supplementary Figure S6c).

**Figure 2:**
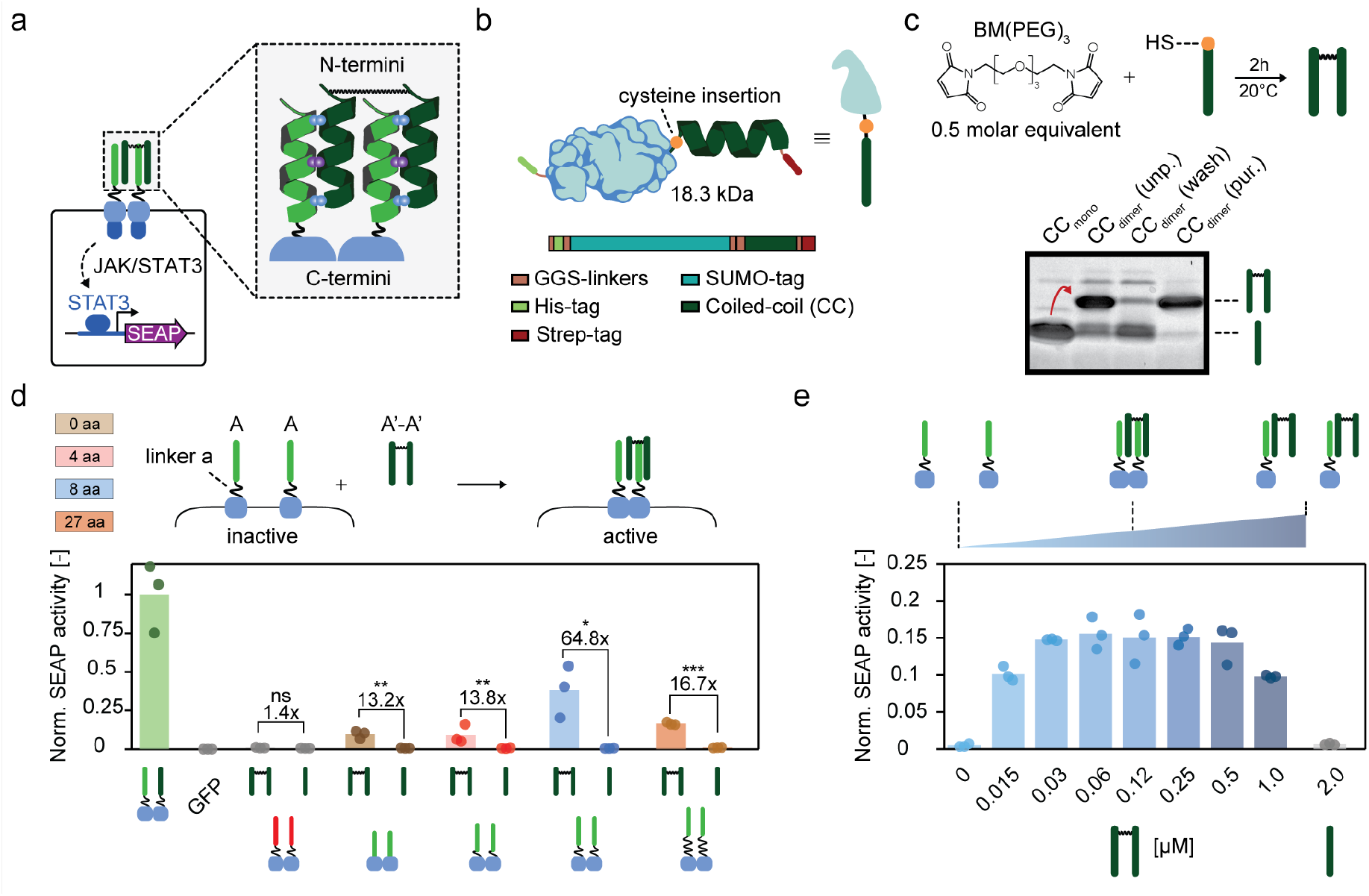
Synthetic, soluble, ditopic CC ligands result in robust receptor activation. **a** Schematic overview of the envisioned ligand-receptor interaction. N-termini linkage of two monomeric CCs results in the desired parallel conformation of a bivalent CC ligand for receptor activation. **b** Schematic representation of the expression strategy of the monomeric CC ligand in *E. coli*. An N-terminus, cleavable small ubiquitin-like modifier (SUMO) tag is fused to a CC ligand, bearing a single cysteine at the C-terminus. For purification purposes, the fusion protein harbours an N-terminal hexahistidine (His) tag and a C-terminal Strep-tag, connected through a small, flexible glycine-serine (GS) linker (see Methods and Supplementary Figure S4). **c** SDS-PAGE analysis showing the successful engineering of the dipeptide prior to purification (CC_dimer_(unp.). Anion-exchange chromatography results in removal of unreacted CC monomers (CC_dimer_(wash) and recovery of the ditopic CC ligands (CC_dimer_(pur.) (see Methods and Supplementary Fig. S4). **d** Normalised SEAP activity in HEK293T cells transiently transfected with CC-GEMS receptors (A-A or B-B) with varied linker lengths **α** of zero, four, eight and 27 aa (GS or GSS repeats). Following transfection, cells were incubated with either 0.12 μM purified A’-A’ dipeptide or 0.24 μM A’ monomeric CC and SEAP expression was monitored to assess receptor activation (see Methods). Data shown are normalised on a heterodimeric A-A’ receptor positive control (green bar). Fold change and significance (unpaired t-test) is noted above bars. ns: p>0.05, *p ≤0.05, **p ≤0.01, ***p ≤0.001 (see Supplementary Table S5). **e** Titration of ditopic ligand A’-A’ on HEK293T cells transiently transfected to express the A-type receptor with **α**_**2**_ linker (eight aa, GS repeats). Cells were incubated with a range of concentrations of ligand A’-A’ (0-1 μM) or 2 μM of monomeric A’ for 48h. SEAP activity was monitored to assess receptor activation. Bars indicate mean activity; individual data points represent independent triplicates.

After successful synthesis and purification of A’-A’ dipeptides, we aimed to evaluate their ability to dimerize and activate A-type receptors. For this purpose, we transfected HEK293T cells with A-type receptors, STAT3 and SEAP reporter genes and incubated the transfected cells with A’-A’ dipeptides or A’ monomeric peptides (A’.C4G (**v**_**1**_); see Methods and Figure 2d). Our results reveal that A’-A’ dipeptides can activate A-type receptors, as evident by an increase in SEAP expression. As expected, no receptor activation was observed for monomeric A’. Both A’ monomers and A’-A’ dipeptides failed to activate non-cognate B-type receptors, as expected. To investigate the influence of linker **α**_**x**_ (see Supplementary Table S2) between the A-type CC and EpoR domain on ligand-dependent receptor activation, we expressed cells with A-type receptors with no linker, linker **α**_**1**_, **α**_**2**_, or **α**_**3**_, and incubated with A’-A’ dipeptide. We observe a 13.2-fold increase for linker **α**_**x**_ of 0 aa, 13.8-fold increase for linker **α**_**1**_, 64.8-fold increase for linker **α**_**2**_ and a 16.7-fold increase for the longer linker **α**_**3**_, when compared to incubation with monomeric A’s. We therefore used receptors with **α**_**2**_ linker in further experimental set-ups. Interestingly, we noticed a decrease of approximately 60% SEAP expression for ligand-induced type activation compared to the situation in which cells express cognate A-type and A’-type receptor pairs with linker **α**_**2**_ (Figure 2d; blue and green bars). We hypothesised that these differences in activation levels can be explained by the number of dimerised CC-GEMS receptors. Specifically, when cognate CC-GEMS are concurrently expressed, dimerization can, in principle, already take place within the secretory pathway before CC-GEMS are translocated to the cell membrane. This would result in increased activation levels, compared to CC-GEMS that are dimerised due to a cognate CC dipeptide ligand, since soluble ligands can only reach receptors that are present on the cell membrane. Ligand-induced receptor dimerization typically results in a bell-shaped dose-response curve that is very susceptible to changes in ligand concentration^50^. When titrating A’-A’ dipeptides to HEK293T cells transiently transfected with A-type receptors with linker **α**_**2**_, STAT3 and SEAP reporter genes, a bell-shaped dose-response curve with a broad plateau, for concentrations ranging between 0.03 and 0.5 μM was observed (Figure 2e).

To further investigate ligand design, we expressed two alternative ditopic A’-A’ ligands (see Supplementary Table S4) that contained a small N-terminal linker that included either one (**v**_**2**_ in Supplementary Figure S6a, S6b and S6d) or two repeats (**v**_**3**_ in Supplementary Figure S6a, S6b and S6e) of the strong helix-interrupting residues proline and glycine^51^. Experiments reveal that the alternative ditopic A’-A’ ligands **v**_**2**_ and **v**_**3**_ result in robust receptor activation in HEK293T cells that are transiently transfected with A-type and reporters (Supplementary Figure S6f and Supplementary Table S6).

These results collectively demonstrate that synthetic, CC ligand dipeptides can activate cognate CC-GEMS receptors.

Having established that synthetic, N-termini linked dipeptides can activate cognate CC-GEMS receptors, resulting in the expression of the reporter gene, we next sought to engineer a ditopic CC ligand that can be expressed and secreted, and can thus facilitate inter-cellular communication. Accordingly, we envisaged two CC peptides that are genetically fused through a long polypeptide linker (**l**), bridging the N-terminus of one CC to the C-terminus of another (see Figure 3a). The anticipated minimum length that a linker needs to bridge between the C-terminus of one CC domain and the N-terminus of another has been estimated to be around approximately 4 nm (calculated distance based on the length of α-helix for a single CC domain, see Methods). To span the two CCs, we made use of three linkers, comprising a combination of rigid α-helical blocks ((EAAAK)_6_)^52^ and more flexible domains ((SGSSGS)_3_)^53^, namely, **l**_**1**_, **l**_**2**_, and **l**_**3**_ (see Methods, Figure 3b and Supplementary Table S7)^54,55^. To engineer A’-A’ dipeptides with linker **l**_**x**_, we initially employed a bacterial expression system (see Methods and Supplementary Figure S5b). Successful expression and purification of the SUMO-tagged A’-A’ was confirmed using SDS-PAGE and successive SUMO cleavage (Methods and Supplementary Figure S7a) resulted in the recovery of pure A’-A’ dipeptides, with linker **l**_**1**_ (Supplementary Figure S7b), **l**_**2**_ (Supplementary Figure S7c) and **l**_**3**_ (Supplementary Figure S7d). To assess the ability of recombinantly expressed A’-A’ ditopic peptides connected with a linker **l** to activate cognate CC-GEMS receptors, we transiently transfected HEK293T cells with A-type receptors with **α**_**2**_, STAT3 and SEAP reporter genes and incubated the transfected cells with the A’-A’ ligands with and without the SUMO fusion (Methods and Figure 3c). We observed reporter gene expression for all the cognate ligand-receptor pairs, while the presence of the SUMO-tag did not inhibit receptor activation (see Supplementary Table S5). A’-A’ ligands with a semi-flexible **l**_**2**_ spacer (Figure 3c; yellow bars) triggered similar receptor activation levels as synthetic A’-A’ ligands (Figure 3c; blue bar), demonstrating 15.9-fold increase for synthetic A’-A’ and 14.5-fold increase for expressed A’-A’ with **l**_**2**_, when compared to cells incubated with A’ monomers. A’-A’ ligands with a shorter **l**_**1**_ linker resulted in even higher receptor activation. More specifically, we observed a 21.3-fold increase for SUMO tagged A’-A’ with **l**_**1**_ and 14.5-fold for A’-A’ with **l**_**1**,_ when compared to cells incubated with A’ monomers. A’-A’ ligands with the longest linker **l**_**3**_ resulted in the lowest receptor activation; 8.7-fold for SUMO tagged A’-A’ with **l**_**3**_ and 7.8**-**fold for A’-A’ with **l**_**3**_, when compared to cells incubated with A’ monomers. Native-PAGE analysis following protein purification (see Methods and Supplementary Figure 8) revealed the presence of a distinct protein band, indicating that A’-A’ dipeptide does not aggregate. To summarize, these results prove the ability of recombinantly expressed soluble, ditopic CC ligands to induce receptor dimerization and subsequent activation.

**Figure 3:**
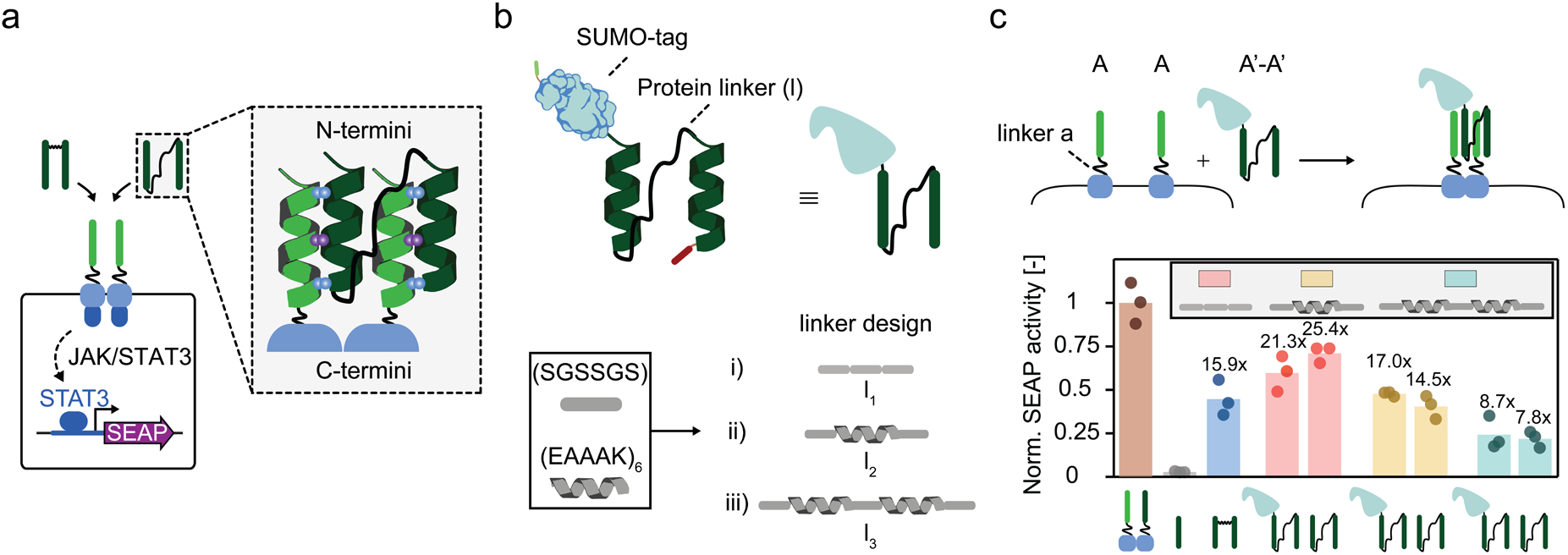
Ditopic CC ligands with various protein linkers activate cognate receptors. **a** Alternative ditopic CC ligands expressed in bacteria are engineered by fusing the N-terminus of a monomeric CC to the C-terminus of another, using various protein linkers. **b** The ditopic ligand was expressed in *E. coli*, after SUMO-tagging. Flexible (SGSSGS) and more rigid (EAAAK) sub-units were used to engineer a total of three linkers (**l**_**x**_)^52,53^ (See Methods, Supplementary table S7 and Supplementary Figure S7) **c** Normalised SEAP activity in HEK293T cells transiently transfected with A type receptor with linker length **α**_**2**_ (8 aa, GS repeats). Following transfection, cells were incubated with either 0.12 μM purified A’-A’ dipeptide ligand, with or without a SUMO-tag, bearing various linker lengths (**l**_**1**_: pink bars, **l**_**2**_: yellow bars, or **l**_**3**_: light blue bars), 0.12 μM A’-A’ reference ligand (ref.: blue bar; see Figure 2c), or 0.24 μM A’ monomeric CC (see Methods). SEAP expression was monitored to assess receptor activation. Data shown are normalised on a heterodimeric A-A’ receptor positive control (brown bar). Bars indicate mean activity; individual data points represent independent triplicates. Fold changes are calculated against cells incubated with A’ monomeric CC and are shown above bars (see Supplementary Table S5).

### Scalability, orthogonality, and two-input logic bio-computations using CC-GEMS

By exploiting the demonstrated ability of designated CCs to specifically and exclusively interact with a cognate receptor partner^40,41,51,56^, we next aimed to explore CC-GEMS receptor activation by an assortment of different dipeptide ligands, thus showing the scalability and programmability of the platform. To this end, CC-GEMS receptors with linker **α**_**2**_, harbouring A, B, or Γ CC domains were engineered (Supplementary Table S1, S2, Supplementary Figure S1) as well as the cognate, recombinant A’-A’, B’-B’ and Γ’-Γ’ dipeptide ligands (see Supplementary Figure S9 and inset in Figure 1a for cognate pairs). To span individual CCs, we opted for the semi-rigid, helical linker **l**_**2**_, described in the previous section. This design choice was made based on the evidence that revealed increased expression of proteins containing helical linker domains in mammalian cells^57–59^. To assess CC-GEMS receptor activation from the designed cognate ligands, HEK293T cells were transfected with A-type, B-type, or Γ-type receptors, as well as STAT3 and SEAP reporter genes and were subsequently incubated with A’-A’, B’-B’ or Γ’-Γ’ dipeptide ligands (Methods). Our results show that only cognate dipeptide ligands are able to activate the corresponding CC-GEMS receptors, while non-cognate pairs result in negligible activation (see Figure 4a and Supplementary Table S6). Interestingly, A’-A’ dipeptide-A-type receptor pairing resulted in the highest activation, followed by Γ’-Γ’ dipeptide-Γ-type receptor pairing and B’-B’ dipeptide-B-type receptor pairing; approximately 30-fold, 16-fold, and fivefold increase respectively. The higher binding affinity of A’:A when compared to B’:B and Γ’:Γ has been previously demonstrated^41^, suggesting that increase in receptor activation might be controlled by the binding affinity of individual CCs. These results indicate that cell response can be rationally designed by linking the predictability of the apparent dissociation constant (*K*_*D, app*_) of each CC module to receptor activation levels. Additionally, incubating A-type receptor expressing cells with an equimolar mixture of A’-A’, B’-B’ and Γ’-Γ’ ligands exhibited similar activation levels to incubation with A’-A’ ligand alone (Supplementary Figure S10); demonstrating that non-cognate CC dipeptides do not inhibit cognate dipeptide-receptor binding. Collectively these data establish CC-GEMS as a scalable, tunable, and orthogonal platform for synthetic receptor activation that can be used for engineering of diffusible, intercellular communication for mammalian cells.

**Figure 4:**
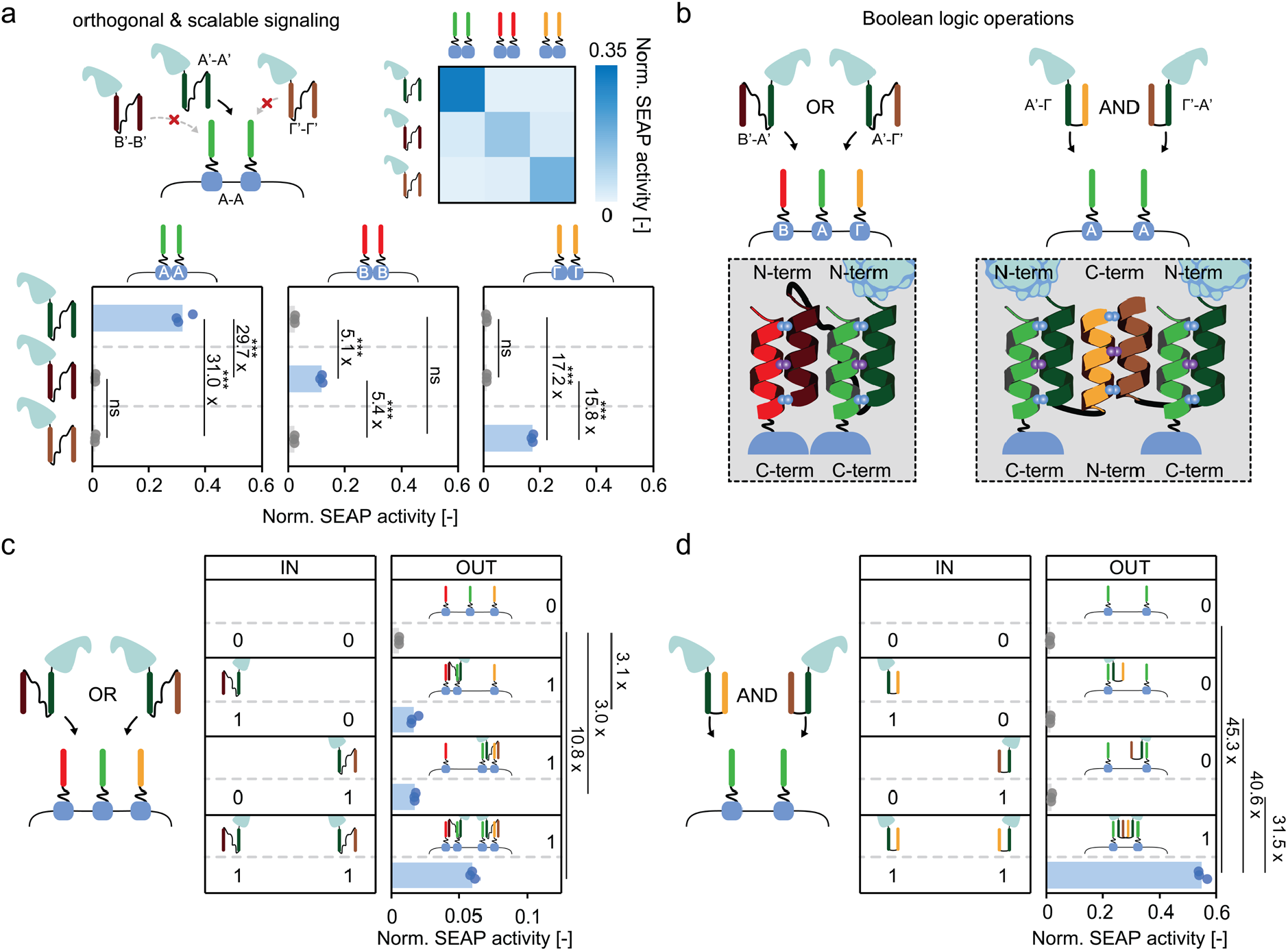
The CC-GEMS platform is scalable and programmable. **a** Normalised SEAP activity in HEK293T cells transiently transfected with CC-GEMS receptors (bottom panel; A-type: left, B-type: middle, Γ-type: right) with linker **α**_**2**_ (8 aa, GS repeats), and subsequently incubated with 0.12 μM purified A’-A’, B’-B’ or Γ’-Γ’ SUMO-tagged dipeptide ligand, with **l**_**2**_ linker (see Methods). The heat-map on the upper right corner depicts the normalised mean SEAP activity per receptor-ligand pair (white: low SEAP activity, blue: high SEAP activity). **b** Schematic representation of envisioned Boolean logic operations. For the OR gate, HEK293T cells are transiently transfected with A-, B- and Γ-type receptors followed by addition of ligands B’-A’ and/or A’-Γ’ to cell culture. For the AND gate, HEK293T cells are transiently transfected with A type receptor and later incubated with A’-Γ and A’-Γ’ ligands. **c** Normalised SEAP activity in HEK293T cells transiently transfected with receptor heterodimers A, B and Γ with linker **α**_**2**_ (8 aa, GS repeats), and afterwards incubated with 0.5 μM SUMO-tagged purified ligand B’-A’ and/or A’-Γ’ with linker **l**_**2**_ (see Methods). Left panel (IN; input) denotes the presence (1) or absence (0) of ditopic ligand. Right panel (OUT; output) shows normalised SEAP activity. **d** Normalised SEAP activity in HEK293T cells transiently transfected with A-type receptors with linker **α**_**2**_ (8 aa, GS repeats), and incubated with 0.12 μM SUMO-tagged purified ligand A’-Γ and/or Γ’-A’ with linker length **l**_**4**_. Left panel (IN; input) denotes the presence (1) or absence (0) of ditopic ligand. Right panel (OUT; output) shows normalised SEAP activity. Data shown are normalised on a heterodimeric A-A’ receptor positive control (not shown). Bars indicate mean activity; individual data points represent independent triplicates. Fold change and significance (one way ANOVA with Tukey’s multiple comparison test) is noted besides the bars. ns: p>0.05, *p ≤0.05, **p ≤0.01, ***p ≤0.001. (see Supplementary Table S6).

The proven orthogonality of cognate CCs^40,41,51,56^ makes CC-GEMS an ideal platform for the implementation of Boolean logic. For instance, engaging an assortment of designed CC-GEMS receptors and dipeptides can enable receptor activation based on combinatorial ligand sensing, as well as OR gate logic operations (Figure 4b). For proof of principle, we expressed two ditopic, SUMO fused B’-A’ and A’-Γ’ ligands (Supplementary Figure S11a and S11b), able to dimerize cognate B-type and A-type as well as A-type and Γ-type receptors respectively (Supplementary Figure S11c). To construct OR gate logic, we transfected HEK293T cells with A-type, B-type, and Γ-type receptors as well as STAT3 and SEAP reporter genes (see Methods) and subsequently incubated them with either B’-A’ or A’-Γ’ ligands with a SUMO-tag, or an equimolar mix of both (Figure 4c). SUMO fused B’-A’ or A’-Γ’ ligands caused an almost three-fold increase in receptor activation, while incubation with both ligands resulted in a 10.8-fold increase. The lower receptor activation due to incubation with either B’-A’ or A’-Γ’ ligands is most likely caused by the fact that expressing three receptor types restricts transfection efficiency and lowers the probability of assembling the appropriate receptor combination at the membrane. Transient transfection with three unique genes results in a heterogenous population of mammalian cells, expressing three, two or one genes^60^, limiting the availability of receptors to cognate ligands. We show that SUMO-tagged B’-A’ dipeptides can result in SEAP excretion in cells that express A and B-type receptors, achieving a 47.4-fold receptor activation. Similarly, SUMO-tagged A’-Γ’ ligand resulted in a 54.0-fold expression of the reporter gene in cells that express A-and Γ-type receptors (Supplementary Figure S11d).

Next, we established combinatorial CC ligand sensing by means of AND gate logic, by expressing A’-Γ and Γ’-A’ dipeptide ligands as a fusion protein with SUMO (Supplementary Figure S12). In this design, Γ:Γ’ pairing facilitates inter-ligand dimerization, giving rise to a CC-CC complex with two accessible A’ domains, available for receptor binding (see Figure 4b; right panel). As Figure 4d illustrates, only in the presence of both SUMO-tagged A’-Γ and Γ’-A’ ligands, receptor activation in HEK293T cells transiently expressing A-type receptor and STAT3 and SEAP reporter genes is observed. In detail, incubation with both A’-Γ and Γ’-A’ resulted in approximately 30-to 45-fold increase compared to incubation with A’-Γ or Γ’-A’ alone or when no ligand was present. In this ligand design, we opted for a shorter linker **l**_**4**_ spanning the individual CCs, assuming that the Γ:Γ’ interaction can act as a natural spacer allowing parallel orientation for the two available A’ CCs, rendering the use of a longer linker unnecessary. Using Native-PAGE we reveal that, following incubation of purified, equimolar SUMO-tagged A’-Γ and Γ’-A ligands the A’-Γ:Γ’-A’ complex forms (see Supplementary Figure S13a and S13b). Size Exclusion Chromatography (SEC) analysis of incubated, equimolar concentrations of SUMO-tagged A’-Γ and A’-Γ’ dipeptides shows that incubation of dipeptides for 24 hours results in complex formation (see Supplementary Figure S13c and S13d). Furthermore, we see that incubation of cells expressing A-type receptor and reporter that are incubated with A’-Γ’ dipeptide do show a significant increase in SEAP production when compared to cells that are not treated with ligand (Supplementary Figure S14 and Supplementary Table S6). However, this 1.4-fold increase of incubation with A’-Γ’ dipeptide, when compared to incubation with no ligand is negligible in comparison to the 45.3-fold activation we see for the AND gate. These results indicate that A’-Γ’ dipeptides can cause some A-type unspecific activation, albeit at minor levels. Additionally, we incubated HEK293T cells expressing A-type receptor and STAT3 and SEAP reporter genes with A’-Γ and Γ’-A’ dipeptide ligands, where the SUMO tag was cleaved and saw robust reporter gene expression, with an approximate 45-55-fold receptor activation (Supplementary Figure S15) suggesting that the presence of a SUMO tag does not inhibit receptor activation. Thus, our data shows that Boolean operations can be engineered at the receptor level utilizing the CC-GEMS platform.

### Establishing intercellular communication using CC-GEMS

Design of intercellular communication between mammalian cells has been limited by the availability of orthogonal signalling mediators. Therefore, we sought to realize a synthetic, intercellular, communication system in mammalian cells that consists of a sender population secreting soluble, bifunctional CC ligands and a receiver cell population expressing the cognate CC-GEMS receptor and reporter genes. To prove that CC dipeptide ligands secreted by mammalian cells can activate CC-GEMS receptors, we stably transfected HEK293T cells to express and subsequently excrete SUMO-tagged A’-A’ dipeptide with linker **l**_**2**_ (Senders; Supplementary Figure S16). Following lentiviral transduction (see Methods), we obtained a heterogenous population, with 84.4% of cells expressing A’-A’ ligand - as quantified by the detection of EmGFP expressed as a bicistron (see Supplementary Figure S16 and S17a). The SUMO-tagged A’-A’ dipeptide was successfully recovered from the medium of sender cells following purification (Supplementary Figure S17c) and it was estimated that sender cells can produce sufficient amounts of A’-A’ ligand to activate receptors on receiver cells (approximately 16.1 nM; see Methods and Supplementary Figure S17d). Next, the mammalian cell-secreted SUMO-tagged A’-A’ purified ligand was added to the medium of HEK293T cells transiently transfected to express A-type receptor as well as STAT3 and SEAP reporter genes. Our data shows that A’-A’ dipeptide ligands expressed in mammalian cells can activate A-type receptors, albeit in a moderately decreased fashion compared to A’-A’ dipeptide ligands isolated from bacteria (Supplementary Figure S17e). To achieve transcriptional control over signalling CC ligand expression, we placed the A’-A’ gene under the control of a doxycycline-induced cytomegalovirus (CMV) promoter^61,62^ (Methods and Supplementary Figure S16) and transduced HEK293S GnTi^-^ TetR cells^61^ (Methods and Supplementary Figure S18a, S18c). Next, cells were enriched for the expression of the transgene (Supplementary Figure S18b) and doxycycline-induced expression was assessed (Supplementary Figure S18d and S18e). The selected population attained 95.3% ligand expression upon induction with doxycycline; however, we observed moderate expression of the transgene in the absence of the inducer in accordance with studies indicating basal leakiness of stably introduced Tet controlled systems^63,64^.

To engineer receiver cells, we first aimed to construct a SEAP reporter cell line by stably transfecting HEK293T cells with the reporter gene (see Methods, Supplementary Figure S19 and S20a). To that end, HEK293T cells were transduced to express SEAP, under the control of four repeats of the STAT3 recognition element and a constitutive mCherry marker (Methods, Supplementary Figure S19, S20a and S20c). Next, a monoclonal cell line population was selected (Supplementary Figure S20b and S20d), which was subsequently transiently transfected with cognate A and A’ type receptors to assess reporter cell line functionality. Figure S20e illustrates that the reporter cells exhibited a 3.7-fold increase in SEAP activity when expressing cognate A- and A’-type receptors compared to cells that expressed A-type receptors alone. Additionally, reporter cells exhibited a 13.8-fold increase in SEAP activity as opposed to untransduced HEK293T cells, indicating “leaky” expression of SEAP at baseline. Receiver cells were engineered by stably transfecting the SEAP reporter cell line with A type receptor constructs with linker **α**_**2**_ (see Methods and Supplementary Figure S21a). When receiver cells were incubated with A’-A’ dipeptide ligand, we observed a 3.7-fold increase in SEAP activity (Supplementary Figure S21b and S21c), indicating that receiver cells are functional and can respond to cognate dipeptide ligands. To demonstrate the ability of the CC-GEMS platform to facilitate intercellular communication, we co-cultured sender and receiver cells (see Methods) in the presence and absence of inducer (Figure 5a). Our data show that intercellular communication can be achieved by co-culturing sender and receiver cells and that reporter gene expression can be temporally controlled with an ON/OFF switch (Figure 5b).

**Figure 5:**
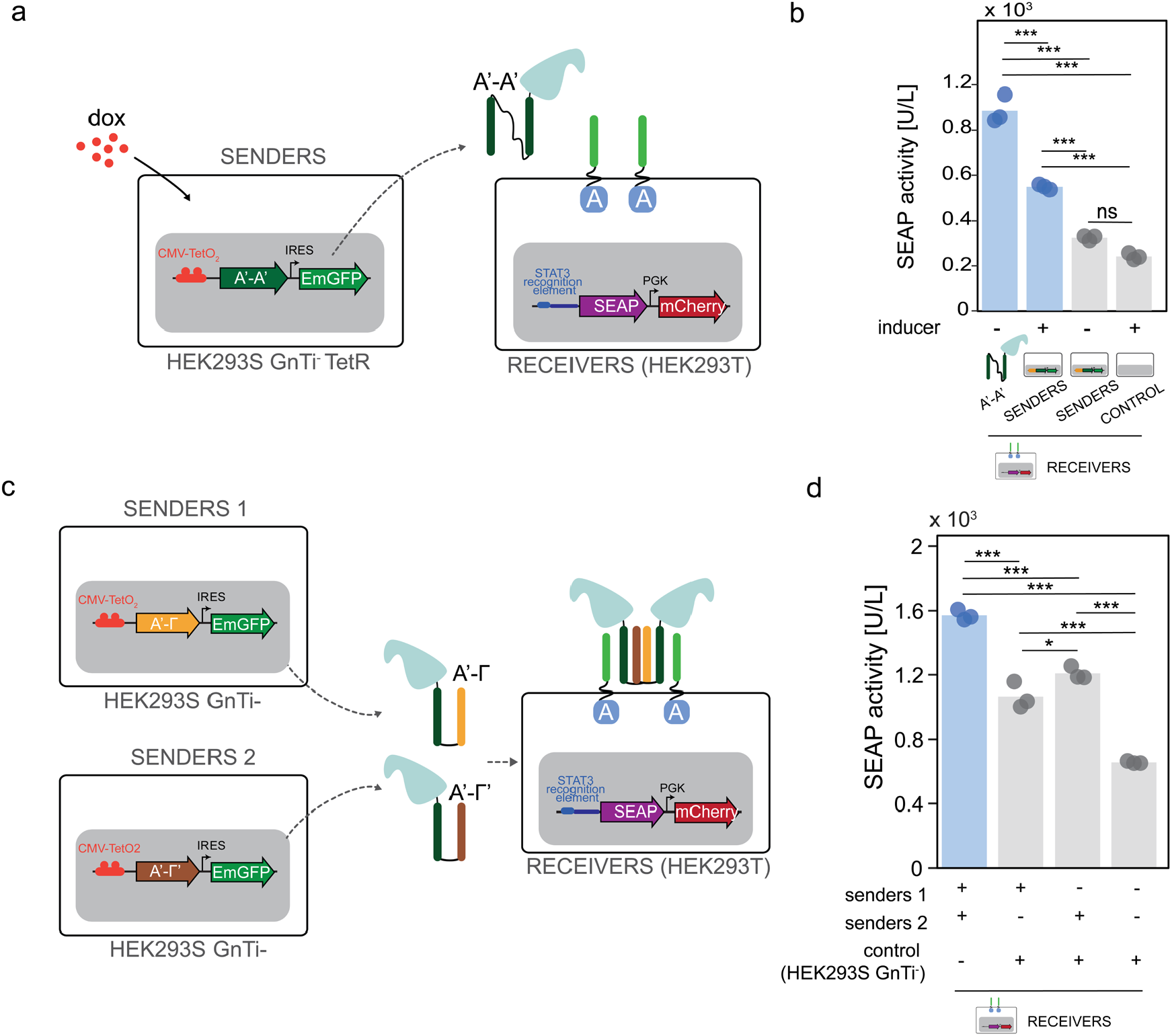
Engineering intercellular communication using CC-GEMS. **a** Intercellular communication is achieved by engineering sender HEK293S GnTi^-^ TetR cells that, upon induction with doxycycline (dox), excrete SUMO-tagged ditopic A’-A’ ligand (see Methods, Supplementary Figure S16 and Supplementary Figure S18) and receiver HEK293T cells expressing the receptor and reporter (see Methods, Supplementary Figure S19, Supplementary Figure S20 and Supplementary Figure S21). **b** SEAP activity (in U/L) in co-cultures of receiver (1.2 × 10^5^ cells) and sender cells (1.2 × 10^5^ cells) or control (untransduced HEK293S GnTi^-^ TetR; 1.2 × 10^5^ cells) following incubation with and without dox (see Methods). As a control, receiver cells were additionally incubated with 0.12 μM ditopic A’-A’ ligand, expressed in *E. coli*, in the absence of sender cells. **c** AND gate logic is achieved by engineering sender 1 (see Supplementary Figure S23) and sender 2 (see Supplementary Figure S24) HEK293S GnTi^-^ cells that secrete SUMO-tagged ditopic A’-Γ and Γ’-A’ ligands respectively (see Methods) and receiver HEK293T cells. **d** SEAP activity (in U/L) for AND gate logic, after incubation of receivers (3.6 × 10^5^ cells) with senders 1, senders 2 and/or control, untransduced HEK293S GnTi^-^ cells (1.2 × 10^5^ cells) for 48 hours in the presence of dox(see Methods). Bars indicate mean activity; individual data points represent independent triplicates. Significance (one way ANOVA with Tukey’s multiple comparison test) is noted above the bars. ns: p>0.05, *p ≤0.05, **p ≤0.01, ***p ≤0.001 (see Supplementary Table S6).

Finally, we aimed to demonstrate that CC-GEMS is a suitable platform for engineering co-cultures that can perform distributed Boolean logic. As a proof of concept, we engineered a prototype, minimal three-cell population system that consists of two senders and a receiver cell line that together execute AND gate logic (Figure 5c). In detail, sender population 1 was created to excrete SUMO-tagged A’-Γ ligand (stable transfection; Methods, Supplementary Figure S22 and S23) and sender population 2 was engineered to excrete SUMO-tagged A’-Γ’ ligand (Methods, Supplementary Figure S22 and S24). As depicted in Figure 5d, receiver cells incubated with both sender populations (senders 1 and 2) show an increase in the reporter gene production when compared to receiver cells incubated with either sender population 1 or sender population 2 and control HEK293S GnTi^-^ cells. Supplementary Figure S25 shows that the AND gate can achieve similar levels of receptor activation as incubation of receiver cells with 0.12 μM SUMO-tagged A’-A’ ditopic ligand. Additionally, we notice that receivers incubated with either sender population 1 or sender population 2 show a moderate increase in receptor activation when compared to receivers incubated with control cells. This is in accordance with the data shown in Supplementary Figure S14, where we observe moderate increase in SEAP production, in HEK293T cells transiently transfected to express A-type receptor and reporters incubated with A’-Γ or A’-Γ’ ligand compared to control (no ligand). This effect of unspecific receptor activation is most likely heightened by the fact that the receiver cell line exhibits leaky expression of the reporter gene (see Supplementary Figure S21). Therefore, we expect that by resolving reporter leakage-related issues, the effect of unspecific receptor activation, when one ligand is present, will be significantly diminished in comparison to receptor activation in the presence of both AND-gate ligands, similar to what we see in Figure 4d and Supplementary Figure S14b. We additionally show that for different ratios of receiver : sender 1 : sender 2 cells, the distributed AND gate results in higher SEAP activity when compared to receiver cells incubated with either sender 1 or sender 2, or control (see Supplementary Figure S26). Collectively our results establish CC-GEMS as a scalable platform for engineering intercellular communication based on soluble, orthogonal ligands with the potential to perform distributed Boolean operations. However, future research should focus on optimizing such intercellular, synthetic communication platforms, resolving issues related to reporter leakage as well as unspecific receptor activation.

## Conclusion

Engineering synthetic communication in mammalian cells has the potential to uncover the fundamental principles of mammalian cell-cell communication circuits as well as provide a basis for the engineering of specialised cell communication networks with custom-defined functionalities. Here we developed CC-GEMS, a modular mammalian synthetic communication platform based on designed, tunable, orthogonal and diffusible ligands that can perform distributed Boolean logic operations. Our strategy is based on the functionalization of synthetic GEMS receptors^10^ with CCs that are engineered to orthogonally bind to a selected partner^40,41^. The inherent orthogonality of CC pairing potentiates the engineering of a cell network that allows for targeted and specific communication between cells. Additionally, the programmable design of large sets of CC heterodimers^42^ would enable the engineering of scalable cell circuit architectures. We demonstrate here that GEMS receptors functionalised with heterodimeric, cognate CC pairs result in robust receptor activation with little influence of the linker spanning the transmembrane domain of the receptor and the CC domain (Figure 1d). The design of a CC dipeptide ligand that can be biologically expressed allows for subsequent receptor activation in an orthogonal manner (see Figure 3 and 4a), enabling the implementation of Boolean operations, in the form of AND and OR gates (see Figure 4b-d). Finally, by engineering a sender-receiver population system (Figure 5), where sender cells can excrete the ligand of choice under the control of a chemical switch or depending on the physiological process, we provide evidence that our system can be used for the engineering of intercellular synthetic communication in mammalian cells.

We envisage that CC-GEMS has several potential applications, including the design of novel cell-based therapeutic approaches. In detail, although autonomously regulated, population control in mammalian cells has been previously achieved, by repurposing the plant hormone auxin^37^, such an approach is not scalable, limiting the response of therapeutic cell populations. We expect that quorum sensing in mammalian cells could be enabled by CC-GEMS and as such afford the engineering of sophisticated systems of therapeutic cells that resemble the human immune system. While CC-mediated interactions have been recently used to engineer cellular assemblies in a helixCAM platform^65^, CC-GEMS could be used to modulate cellular response depending on cell-cell interactions through membrane presented ditopic CC signalling mediators.

One of the prominent features of CCs is their designable orthogonality and tunability, which allows selection of CCs with appropriate affinity^66^ to obtain the appropriate response under the selected conditions. Introduction of combinations of several dipartite CCs and use of 4-helix bundles could enable combinations of more than 2 input signals and more versatile complex information processing. CC mediators also facilitate construction of biological systems that can respond within minutes as they do not require transcriptional regulation due to their dependence on protein interactions and modifications^67^.

Future research should focus on further developing the inter-cellular communication in co-culture of cells, resolving issues related to gene expression leakage. The reporter gene could be straightforwardly exchanged for fluorescent protein molecules or more relevant therapeutic protein genes. Additionally, engineering primary immune cells with CC-GEMS has the potential to show the ability of our platform to control responses in cell-based therapeutic systems.

## Methods

### Chemicals and reagents

All reagents and solvents were obtained from commercial sources and were used without further purification.

### Gene construct design and preparation

Plasmids were constructed using the appropriate backbones for either transient or stable expression of the transgene (see Table S2) and desired DNA (purchased from Integrated DNA Technologies) with the use of restriction-digestion, with standard restriction enzymes (New England BioLabs; HF enzymes were used when possible) and ligation with T4 DNA ligase (New England BioLabs). All PCRs were performed with Q5 High-Fidelity DNA Polymerase (New England BioLabs) or Phusion High-Fidelity DNA Polymerase (New England BioLabs) according to the manufacturer’s instructions. All plasmids used in this study are listed in Supplementary Table S9. Sequence verification of genes and constructs was done by Sanger sequencing (BaseClear).

### Cell culture and transient transfection

Human embryonic kidney (HEK293T) cells (validated and mycoplasm-free; ATCC® CRL3216™) were maintained in growth medium, Dulbecco’s Modified Eagle Medium (DMEM, 41966; Thermo Fisher Scientific) supplemented with 10% (v/v) Fetal Bovine Serum (FBS, S-FBS-SA-015; Serana) and 1% (v/v) antibiotic penicillin/streptomycin (pen/strep, 10,000 U/mL; Gibco), under standard incubation conditions (37°C, in a humidified atmosphere of 5% CO_2_).

To prepare for transfection, cells were seeded at a density of 2.4×10^5^ cells per well in a 24-well plate (Greiner) containing growth medium. The next day, cells were transfected with 500 ng plasmid DNA (193.8 ng per receptor dimer for a total of 387.6 ng, 96.1 ng STAT3-induced secreted alkaline phosphatase (SEAP) reporter plasmid pLS13 and 16.3 ng STAT3 expression vector pLS15; see Supplementary Table S2) for five hours, using 1.25 μl of the transfection reagent lipofectamine 2000 (Life Technologies) in a total of 200 μl Opti-MEM™ I reduced serum medium (Thermo Fisher Scientific), according to manufacturer’s instructions. Following transfection, cells were incubated with growth medium, containing the appropriate ligand for 48 hours; at 37°C for CCs A and A’ (Figure 1, 2 and 3; Supplementary Figure S3, S6 and S9) and 30°C (Figure 4; Supplementary Figure S9 and S10). Only in the case of the OR gate (Figure 4c), a total of 693.8 ng DNA was transfected (193.8 ng per receptor dimer (3x) for a total of 387.6 ng, 96.1 ng STAT3-induced SEAP reporter plasmid pLS13 and 16.3 ng STAT3 expression vector pLS15). When enhanced green fluorescent protein (EGFP) was used as a transfection control, 2.4×10^5^ HEK293T cells seeded in a 24-well plate were transfected with 500 ng of pHR-EGFPligand plasmid DNA (see Supplementary table S9) as described above and transfection efficiency was assessed with flow cytometry.

### SEAP quantification

SEAP concentration in cell culture medium was quantified in terms of absorbance increase due to para-nitrophenyl phosphate (pNPP), using a commercially available SEAP assay kit (NBP2-25285, Novus Biologicals). Briefly, cell culture medium was harvested, and cell debris was removed by means of centrifugation. Subsequently the medium was heat inactivated at 65°C for 30 min. To measure SEAP activity, cell culture medium was added to SEAP sample buffer and SEAP substrate in a transparent 96 well plate (Thermo Fisher Scientific), according to manufacturer’s instructions (in a final concentration of 0.83 mg/mL pNPP). Absorbance values were measured at 405 nm, at a controlled temperature of 25°C, for 60 min, using a Tecan Spark 10M plate reader (Tecan). To determine sample SEAP concentration, a calibration curve was constructed by titrating known concentrations of the hydrolysed pNPP product para-nitrophenol (pNP) (Supplementary Figure S1). Absorbance units are converted to amount of substrate conversion and SEAP activity is calculated from the slope of the time trace and expressed in units per litre (U/L). One unit is defined as the amount of enzyme that converts 1 μmole pNPP in 5 μL cell culture medium in 1 min at 25°C.

### Recombinant expression and purification of CC peptide ligands

A pET28a(+) vector (Supplementary Table S2) encoding the A’ monomeric CC fused to SUMO tag (Supplementary Figure S5 and Supplementary Table S1) was transformed into *E. coli* BL21(DE3) (Novagen). A single colony of freshly transformed cells was cultured at 37°C, in 500 mL 2xYT medium supplemented with 50 μg/mL kanamycin (Merck). When the OD600 of the culture reached ~0.6, protein expression was induced by addition of β-D-1-thiogalactopyranoside (IPTG, AppliChem), in a final concentration of 1 mM. The induced protein expression was carried out at 25°C for ~18 h and the cells were harvested by centrifugation at 10,000 x g at 4°C for 10 min. The pelleted cells were subsequently resuspended in BugBuster (5 mL/g pellet, Merck) supplemented with benzonase (5 μL/g pellet, Merck) and incubated for 15 min on a shaking table. The suspension was centrifuged at 40,000 x g at 4°C for 30 min and the supernatant was subjected to Ni^2+^ affinity chromatography on a gravity column. Briefly, after the supernatant was loaded on the column, the column was washed with washing buffer (1x PBS, 370 mM NaCl, 10% (v/v) glycerol, 20 mM imidazole, pH 7.4) and the protein was eluted with His-elution buffer (1x PBS, 370 mM NaCl, 10% (v/v) glycerol, 250 mM imidazole, pH 7.4). Cleavage of the N-terminal His-SUMO tag was performed by adding SUMO protease dtUD1 (1:500) to the elution fraction, while dialyzing (molecular weight cut-off (MWCO) 3.5 kDa, Thermo Fisher Scientific) against 2 L of dialysis buffer (50 mM Tris, 50 mM NaCl, pH 8.0) at room temperature for 16 h. The concentrate was applied to a Ni^2+^ column and the flow through, containing the CC ligand, was recovered, snap-frozen and stored at 80°C for subsequent use. When CC monomers were used for synthesis of ditopic CC ligands, a final concentration of 2 mM TCEP reducing agent was added to the protein aliquots. Protein concentration was calculated based on absorption at 280 nm in a Nanodrop 1000 spectrophotometer (ND-1000, Thermo Fisher Scientific), assuming an extinction coefficient calculated using the ProtParam tool on the ExPASy server. Protein purity was assessed on reducing SDS-PAGE.

Similarly, the various ditopic CC ligands with linker length **l**_**x**_, were expressed in *E. coli* BL21(DE3) (Novagen). Various linkers **l**_**x**_ were designed from a combination of (EAAAK)_6_^52^ and (SGSSGS)_3_ ^53^ monoblocks (see Supplementary Table S7). The calculated average distance of a single CC domain was approximated to be 4 nm, based on the 0.15 nm per residue length of an a-helix.

### Synthesis and purification of ditopic A’-A’ CC ligands

Monomeric A’ CC peptide, expressed in *E. coli* as described above, was buffered exchanged to 100 mM Sodium Phosphate, 25 mM TCEP, pH 7 using repetitive washing and centrifugation with an Amicon 3 kDa MWCO centrifugal filter (Merck Millipore). For the synthesis of ditopic A’-A’ CC ligand, 0.5 molar equivalent of homo-bifunctional bismaleimide linker (BS(PEG)_3_; Thermo Fisher Scientific) was added to the protein solution and the reactants were incubated at 20°C, 850 rpm for 3 h. Ditopic A’-A’ CC ligands were purified using anion-exchange chromatography. Briefly, following equilibration of the anion-exchange column (0.5 mL strong anion-exchange spin column; 90010, Thermo Scientific) with purification buffer (50 mM Tris-HCl, pH 8.0), the protein mixture was loaded in 400 μL fractions, according to manufacturer’s instructions. Unreacted monomeric A’ CCs were first eluted by treating the column with purification buffer supplemented with 200 mM NaCl and ditopic A’-A’ CCs were eluted by treating the column with purification buffer supplemented with 300 mM NaCl. Purity of the recovered ditopic A’-A’ CCs was assessed using SDS-PAGE under non-reducing conditions and purified products were snap-frozen and stored at −80°C.

### Lentivirus production

Lentivirus was produced by co-transfecting HEK293T cells (ATCC® CRL3216™) that have reached ~80-90% confluency with the second generation pHR plasmid carrying the desired transgene (Supplementary Table S2) and the vectors encoding for the packaging proteins (pCMVR8.74; gift from Didier Trono (Addgene plasmid # 22036; http://n2t.net/addgene:22036; RRID:Addgene_22036)) and VSV-G envelope (pMD2.G, gift from Didier Trono (Addgene plasmid # 12259; http://n2t.net/addgene:12259; RRID:Addgene_12259)) at a ratio of 1:2:1.2, using the FuGENE® HD transfection reagent (Promega) in Opti-MEM™ I reduced serum medium (Thermo Fisher Scientific). The following day, the medium was refreshed to DMEM (41966; Thermo Fisher Scientific) supplemented with 2% (v/v) FBS (S-FBS-SA-015; Serana) and cells were incubated under standard incubation conditions (37°C, in a humidified atmosphere of 5% CO_2_). 48 h later, the supernatant was harvested, and filtered with a 0.45 μm syringe filter (Merck). The produced lentivirus was pelleted by centrifugation at 50,000 x g, 4°C for two hours, resuspended in DMEM (41966; Thermo Fisher Scientific) supplemented with 2% (v/v) FBS (S-FBS-SA-015; Serana) and either used directly for cell line transduction or snap-frozen and stored at −80°C.

### Sender cell line engineering

HEK293T cells (ATCC® CRL3216™) or human embryonic kidney 293S lacking *N*-acetylglucosaminyltransferase I and expressing the Tet repressor protein (HEK293S GnTI^-^ TetR; obtained from the group of N. Callewaert at the VIB-UGent Center for Medical Biotechnology)^68^ were plated in 24-well plates (Greiner) that upon reaching 10-20% confluency were transduced with lentivirus harbouring the SUMO-tagged A’-A’ peptide with linker **l**_**2**_ (Supplementary Figure S18) or SUMO-tagged A’-Γ or Γ’-A’ peptide with linker **l**_**4**_ (Supplementary Figure S23 and S24), under the control of the major immediate–early (MIE) human cytomegalovirus (CMV) enhancer and promoter and two *TetO* operator sequences^69^ followed by an internal ribosome entry site (IRES) from encephalomyocarditis virus (EMCV) controlling the expression of an emerald green fluorescent protein (EmGFP). In total, 50-100 μL lentivirus was added to the cells, containing transduction medium (DMEM (41966; Thermo Fisher Scientific) supplemented with 10% (v/v) FBS (S-FBS-SA-015; Serana)). Virus-containing transduction medium was replaced by growth media 48-72 h post infection. Cells were bulk-sorted for EmGFP by fluorescence-activated cell sorting (FACS). Cells were cultured under standard incubation conditions (37°C, in a humidified atmosphere of 5% CO_2_) and passaged upon reaching confluency (approximately every 2 days). HEK293S GnTI^-^ TetR cells were routinely cultured in growth medium with the addition of 1 μg/mL blasticidine S HCl (11583677, Gibco) for selection of the TetR trait.

### Mammalian protein expression and purification

For A’-A’ CC protein expression and quantification (see Supplementary Figure S14c and S14d), 3.72 × 10^6^ HEK293T A’-A’ sender cells (Supplementary Figure S14a and S14b) were seeded in a T175 cell culture flask (Greiner), containing 35 mL DMEM (41966; Thermo Fisher Scientific) supplemented with 2% (v/v) FBS (S-FBS-SA-015; Serana) and 1% (v/v) antibiotic penicillin/streptomycin (pen/strep, 10,000 U/mL; Gibco). The cells were cultured at 37°C, in a humidified atmosphere of 5% CO_2_ for four days and the medium was harvested and cell debris was removed by centrifugation at 10,000 x g for 10 min. Subsequently, medium was filtered with a 0.45 μm syringe filter (Merck) prior to loading to a Strep-tag affinity chromatography column. Gravity flow columns (2 mL) were prepared with 400 μL 50% Strep-Tactin® XT Superflow High Capacity suspension (IBA Lifesciences). Protein purification was performed according to manufacturer’s protocol. Washing was conducted with Buffer W (100 mM Tris, 150 mM NaCl, 1 mM EDTA, pH 8.0) and elution with Buffer BE (100 mM Tris, 150 mM NaCl, 1 mM EDTA, 50 mM D-biotin). Purified protein was concentrated with the use of a 3K Amicon® Ultra 0.5 mL Centrifugal filter (Merck), according to manufacturer’s instructions. Proteins were buffered-exchanged in buffer containing 50 mM Tris, 50 mM NaCl, pH 8.0.

For A’-Γ and Γ’-A’ CC protein expression (see Supplementary Figure S19 and S20 respectively), supernatant was harvested from 293S GnTi^-^ TetR sender A’-C (senders 1) and 293S GnTi^-^ TetR sender Γ’-A’ (senders 2) cells that have been continuously cultured in DMEM (41966; Thermo Fisher Scientific) supplemented with 10% (v/v) FBS (S-FBS-SA-015; Serana), 1% (v/v) antibiotic penicillin/streptomycin (pen/strep, 10,000 U/mL; Gibco) and 1 μg/mL doxycycline (dox; D3447, Merck). Strep-tag protein purification was conducted as described above.

### SDS-PAGE and Native PAGE

For SDS-PAGE analysis, 4-20% SDS-PAGE Mini-PROTEAN® TGX Precast gels (Bio-Rad) were used with running buffer (25 mM Tris, 192 mM glycine, 0.1% SDS, pH 8.3; Bio-Rad). Samples were added to SDS sample buffer (62.5 mM Tris–HCl, pH 6.8, 2% SDS, 25% (v/v) glycerol, 0.01% bromophenol blue, and 100 mM DTT) and denatured at 95 °C for 5 min. For Native PAGE analysis, purified ligands were added to Native PAGE sample buffer (75 mM Tris, 576 mM glycine, 30% (v/v) glycerol, 0.01% bromophenol blue) and run in a 4-20% SDS-PAGE Mini-PROTEAN® TGX Precast gels (Bio-Rad) with running buffer (25 mM Tris, 192 mM glycine). To assess A’-Γ:Γ’-A’ complex formation, equimolar concentrations of purified A’-Γ and Γ’-A’ ligands were first incubated at 30°C for 1 hour and the formed complex was run in a Native PAGE gel (Supplementary Figure S13). Protein visualization was done with Coomassie Brilliant Blue G-250 (Bio-Rad). Gels were analysed with the ImageQuant 350 (GE Healthcare) or Amersham ImageQuant 800 (Cytiva). All procedures were carried out according to manufacturers’ protocols.

### Size Exclusion Chromatography

Equimolar concentrations of SUMO-tagged A’-Γ and A’-Γ’ dipeptides with protein linker **l**_**4**_ were incubated for one hour or 24 hours and were subsequently characterised using SEC (see Supplementary Figure S13c). The analytical SEC data reported was performed on a NGC Chromatography System (Bio-Rad) using a Superdex 200 Increase 10/300 GL column (GE Healtchare). 1x PBS was used as the running buffer.

### Gel densitometric analysis

Background-subtracted density of the SDS-PAGE gel protein bands was determined as a function of area under the curve of intensity profile plots, using the gel analysis plugin of ImageJ 1.53a. Optical densities (in a.u.) of protein A’-A’ CC ligand expressed in *E. coli* with known concentrations (namely 0.5, 1, 1.5 and 2 μM, Supplementary Figure S17d) were measured and linearly fitted to construct a calibration curve that was later used to determine A’-A’ CC dipeptide concentration expressed by mammalian cells.

### Receiver cell line engineering

For the engineering of the reporter cell lines, HEK293T cells (ATCC® CRL3216™) were plated in 24-well plates (Greiner). Upon reaching 10-20% confluency, cells were transduced with lentivirus so as to express the SEAP reporter gene, under the control of a STAT3 recognition element, which consists off a minimal cytomegalovirus promoter (minCMV) and four STAT3 binding sites. For fluorescent cell tracking, the same construct encodes for mCherry fluorescent protein, under the control of a constitutive phosphoglycerate kinase (PGK) promoter (Supplementary Figure S15 and Supplementary Figure S16). As described above, 50-100 μL lentivirus was added to the cells, containing transduction medium (DMEM (41966; Thermo Fisher Scientific) supplemented with 10% (v/v) FBS (S-FBS-SA-015; Serana). Virus-containing transduction medium was replaced by growth media 48-72 h post infection. Cells were sorted as single cells, in flat-bottom 96-well plates (Greiner) based on mCherry expression, using fluorescence-activated cell sorting (FACS). Following 14 days of seeding, wells were monitored for monoclonal cell line growth. To engineer receiver cell lines, a reporter cell line culture was selected (Supplementary Figure S16e) and was lentivirally transduced with the gene encoding for the A-type receptor with linker **α**_**2**_ (8 GS repeats), as previously described. Cells were cultured under standard incubation conditions (37°C, in a humidified atmosphere of 5% CO_2_) and passaged upon reaching confluency (approximately every 2 days).

### Sender-Receiver co-culture

A total of 1.2 × 10^5^ receiver cells and 1.2 × 10^5^ A’-A’ sender cells (Figure 5b) cells per well were seeded in a 24-well plate (Greiner) in growth medium. For the AND gate logic (Figure 5d), a total of 3.6 × 10^5^ receiver cells and 1.2 × 10^5^ sender 1 and/or 1.2 × 10^5^ sender 2 cells or 1.2 × 10^5^ control (untransduced HEK293S GnTi^-^) cells were co-cultured in a 24-well plate (Greiner) in growth medium. For the AND gate logic experiments presented in Supplementary Figure S26, different ratios of receivers : senders 1 : senders 2 were seeded in a 24 well plate (Greiner) in growth medium. The exact ratios and number of cells seeded are reported in Supplementary Table S10. The populations were incubated in 300 μL growth medium with or without the addition of 1 μg/mL dox for 48 h, at 37°C (Figure 5b) and at 30°C (Figure 5d and Supplementary Figure S26); in a humidified atmosphere of 5% CO_2_. The cell culture medium was subsequently used to determine concentration of SEAP as described above.

### Flow cytometry analysis and fluorescence-activated cell sorting

Cells were interrogated and sorted on a FacsAria III (BD Biosciences), operating at low-middle pressure, using a 70 μm nozzle. Cells were interrogated using a 488 nm laser and a 530/30 nm detector for EmGFP and a 561 nm laser and a 610/20 nm detector mCherry. A total of 10,000-100,000 events were recorded, from which 2D plots of the side-scattered light area (SSC-A) versus forward-scattered light area (FSC-A), as well as FSC-A versus forward-scattered light height (FSC-H) were obtained.

### Data analysis and statistics

Data were plotted with the use of MATLAB Toolbox Release R2019a for Windows (The MathWorks, Inc., Natick, Massachusetts, United States) and show the mean as a bar diagram overlaid with a dot plot of individual data points (for n=3 biologically independent samples). Statistical analysis was done with GraphPad Prism 9.4.1 for Windows (GraphPad Software, San Diego, California USA, www.graphpad.com). All analyses were executed using unpaired, two-sided t-test in case of comparing two experimental groups or One-way Analysis of Variance (ANOVA) with Tukey’s multiple comparisons test, when comparing three or more experimental groups, Bonferroni test (for planned comparisons) or Dunnett’s test (when comparing means to a control group). Only values of p<0.05 were considered statistically significant. Flow cytometry data were analysed with the use of FlowJo™ v10.7.1 Software (BD Life Sciences) for Windows.

## Data availability

The datasets generated during and/or analysed during the current study are available from the corresponding author upon request.

## Author Contributions

A.M.P and G.A.O.C designed the study, performed experiments, and analysed the data. A.M.P wrote the manuscript. A.M.P and B.L.N established cell lines and performed related experiments. B.W.A.B, G.A.O.C and A.M.P analysed and plotted all SEAP quantification experiments. B.V.E and M.T.H.B assisted in cloning and cell experiments. A.d.D. performed experiments related to protein characterization. O.M.S and C.C.V.B established the lentiviral production and provided key insights with regards to stable transfections. M.M, W.J.M.M, and R.J provided critical feedback on experiments and revised the manuscript. T.F.A.d.G conceived, designed and supervised the study, analysed the data and wrote the manuscript.

## Acknowledgements

The authors thank R. Driessen for useful discussions regarding stable transfections, and J. Lentjes for her help with protein expression. This work was supported by the European Research Council (ERC project no. 101000199 AMIGA) grant.

