## Supplementary Information for "Engineering a Scalable and Orthogonal Platform for Synthetic Communication in Mammalian Cells"

### Supplementary Tables

**Supplementary Table S1: DNA and protein sequences of unique monomeric CC modules.** Overview of nucleotide and amino acid sequences of monomeric CCs (A, A’, B, B’, Γ, and Γ’).

| ID | DNA Sequence (5’ to 3’) | Amino acid sequence (N-term to C-term) |
|---|---|---|
| **A** | GAGATCCAGCAACTGGAAGAGGAAATCGCCCAGCTGGAGCAGAAAAACGCCGCTCTGAAGGAAAAGAACCAGGCCCTGAAGTAC | EIQQLEEEIAQLEQKNAALKEKNQALKY |
| **A’** | AAAATTGCGCAGTTGAAACAAAAAATTCAGGCGCTCAAGCAAGAAAATCAACAGTTGGAGGAGGAGAATGCAGCCCTGGAGTAT | KIAQLKQKIQALKQENQQLEEENAALEY |
| **B** | GAGAACGCCGCTCTGGAAGAGAAGATCGCCCAGCTGAAACAGAAGAACGCCGCCCTGAAGGAAGAGATCCAGGCCCTGGAATAC | ENAALEEKIAQLKQKNAALKEEIQALEY |
| **B’** | AAGAACGCCGCTCTGAAGGAAGAGATCCAGGCGTTGGAAGAGGAGAACCAGGCCCTGGAAGAGAAGATCGCCCAGCTGAAATAC | KNAALKEEIQALEEENQALEEKIAQLKY |
| **Γ** | GAGATCCAAGCCCTGGAAGAGAAGAACGCCCAGCTGAAACAGGAGATCGCCGCTCTGGAAGAAAAGAACCAGGCCCTGAAGTAC | EIQALEEKNAQLKQEIAALEEKNQALKY |
| **Γ’** | AAGATCGCCCAGCTGAAGGAAGAGAACCAGCAACTGGAACAGAAGATCCAGGCCCTGAAAGAGGAAAACGCCGCTCTGGAGTAC | KIAQLKEENQQLEQKIQALKEENAALEY |

**Supplementary Table S2: DNA and protein sequences for linker $\alpha_x$.** Three different protein linkers ($\alpha_x$) used to span the CC and EpoR for the construction of the CC-GEMS receptor.

| ID | DNA Sequence (5’ to 3’) | Amino acid sequence (N-term to C-term) |
|---|---|---|
| $\alpha_1$ | GGATCAGGGTCTGGAAGTGGGAGT | GSGSGSGS |
| $\alpha_2$ | GGATCAGGGTCTGGAAGTGGGAGTGGATCAGGGTCTGGAAGTGGGAGT | GSGSGSGSGSGSGSGS |
| $\alpha_3$ | AGTGGATCCTCGGGATCATCAGGATCCTCTGGGAGCTCAGGATCCTCGGGGAGCTCTGGATCCAGCGGTTCAAGCGGATCC | SGSSGSSGSSGSSGSSGSSGSSGSSGS |

**Supplementary Table S3: Plasmid backbones used in the present study.** Overview of plasmid backbones used and corresponding source. Multiple unique plasmids were designed by recombination of unique CC modules and linkers.

| ID | Description | Source |
|---|---|---|
| pLS13 | Mammalian reporter plasmid for STAT3-induced SEAP expression ($O_{Stat3}$-$P_{hCMVmin}$-SEAP-pA). | Schukur et al.[1] |
| pLS15 | Mammalian STAT3 expression vector ($P_{hCMV}$-STAT3-pA). | Schukur et al.[1] |
| pLeo619-$P_{SV40}$ | Mammalian pLeo619 expression vector to recombinantly express the $EpoR_m$-IL-$6RB_m$-pA fused to a selected CC module. | Original plasmid was kindly provided by Martin Fussenenger (GenBank accession no. MG437012)[2]. |
| pET28a(+)-$P_{T7}$ | Bacterial Pet28a(+) expression vector to recombinantly express different variants of monomeric or ditopic SUMO-CC fusion proteins in *E. Coli*. | |

| | | |
|---|---|---|
| pHR-P$_{CMV}$ | Second generation lentiviral transfer pHR-CMV expression vector for generation of lentiviruses. | Gift from A. Radu Arocescu (Addgene plasmid #113891; http://n2t.net/addgene:113891; RRID:Addgene_113891)[3]. |

**Supplementary Table S4: DNA and protein sequences for different modifications on monomeric A' CC module.** Three variants (**v**) of the linker sequence between the SUMO-tag and monomeric A' CC module used to engineer ditopic A'-A' CCs (see Supplementary Fig. S6). The differences with respect to the reference sequence (ref.) are indicated in red.

| ID | DNA Sequence (5' to 3') | Amino acid sequence (N-term to C-term) |
|---|---|---|
| Ref. | AGTGGTTCATGTAGCGGTAGTGGATCC | SGSCSGSGS |
| **v1** | AGTGGTTCAGGTAGCGGTAGTGGATCC | SGSGSGSGS |
| **v2** | AGTGGTTCATGTAGCGGTCCTGGATCC | SGSCSGPGS |
| **v3** | AGTGGTTCATGTAGCGGTCCTGGATCCGGTAGTGGTCCTGGAAGT | SGSCSGPGSGSGPGS |

**Supplementary Table S5: P-values, t-values and degrees of freedom for unpaired t-tests.** Overview of fold change, p-values, t-values and degrees of freedom (df) for unpaired, two-sided t-tests, indicating the compared groups and corresponding Figure. ns: (non-significant), *p ≤0.05, **p ≤0.01, *** p ≤0.001.

| Figure | Compared groups | Fold change | p-value summary | p-value | t-value | df |
|---|---|---|---|---|---|---|
| 2d | B-GEMS with A'-A' vs. B-GEMS with A' | 1.4 | ns | 0.1084 | 2.061 | 4 |
| 2d | A-GEMS (**$\alpha$**=0 aa) with A'-A' vs. A-GEMS (**$\alpha$**=0 aa) with A' | 13.2 | ** | 0.0038 | 6.039 | 4 |
| 2d | A-GEMS (**$\alpha_1$**=4 aa) with A'-A' vs. A-GEMS (**$\alpha_1$**=4 aa) with A' | 13.8 | ** | 0.0733 | 2.413 | 4 |
| 2d | A-GEMS (**$\alpha_2$**=8 aa) with A'-A' vs. A-GEMS (**$\alpha_2$**=8 aa) with A' | 64.8 | * | 0.0183 | 3.850 | 4 |
| 2d | A-GEMS (**$\alpha_3$**=27 aa) with A'-A' vs. A-GEMS (**$\alpha_3$**=27 aa) with A' | 16.7 | *** | <0.0001 | 29.72 | 4 |
| 3c | A-GEMS with SUMO.A'-A' (with **$l_1$**) vs. A-GEMS with A'-A' (with **$l_1$**) | 1.19 | ns | 0.1577 | 1.735 | 4 |
| 3c | A-GEMS with SUMO.A'-A' (with **$l_2$**) vs. A-GEMS with A'-A' (with **$l_2$**) | 1.18 | ns | 0.1409 | 1.832 | 4 |
| 3c | A-GEMS with SUMO.A'-A' (with **$l_3$**) vs. A-GEMS with A'-A' (with **$l_3$**) | 1.11 | ns | 0.7165 | 0.3898 | 4 |
| S21c | receivers with A'-A' vs. receivers without A'-A' | 3.7 | *** | <0.0001 | 31.18 | 4 |

**Supplementary Table S6: P-values, F-values and degrees of freedom for One-Way ANOVA.** Overview of p-values, F-values and degrees of freedom (df) for unpaired, one-way ANOVA with Tukey's, Bonferroni's or

Dunnett's multiple comparison test, indicating the compared groups and corresponding Figure. n.s.: (non-significant), *p ≤0.05, **p ≤0.01, *** p ≤0.001.

| Figure | Compared groups (ANOVA) | F-value | p-value summary (ANOVA) | Compared groups (multiple comparisons) | adjusted p-value (multiple comparisons) | df |
|---|---|---|---|---|---|---|
| S6f | B-GEMS with A'-A' vs. B-GEMS with A'-A' ($\mathbf{v_2}$) vs. B-GEMS with A'-A' ($\mathbf{v_3}$) | 2.724 | ns (p=0.1439) | - | - | 8 |
| S6f | A-GEMS ($\boldsymbol{\alpha}$=0 aa) with A'-A' vs. A-GEMS ($\boldsymbol{\alpha}$=0 aa) with A'-A' ($\mathbf{v_2}$) vs. A-GEMS ($\boldsymbol{\alpha}$=0 aa) with A'-A' ($\mathbf{v_3}$) | 20.73 | ** (p=0.002) | A-GEMS ($\boldsymbol{\alpha}$=0 aa) with A'-A' vs. A-GEMS ($\boldsymbol{\alpha}$=0 aa) with A'-A' ($\mathbf{v_2}$) | ns (p=0.2808) Tukey | 8 |
| | | | | A-GEMS ($\boldsymbol{\alpha}$=0 aa) with A'-A' vs. A-GEMS ($\boldsymbol{\alpha}$=0 aa) with A'-A' ($\mathbf{v_3}$) | ** (p=0.0019) Tukey | |
| | | | | A-GEMS ($\boldsymbol{\alpha}$=0 aa) with A'-A' ($\mathbf{v_2}$) vs. A-GEMS ($\boldsymbol{\alpha}$=0 aa) with A'-A' ($\mathbf{v_3}$) | ** (p=0.0095) Tukey | |
| S6f | A-GEMS ($\boldsymbol{\alpha_1}$=4 aa) with A'-A' vs. A-GEMS ($\boldsymbol{\alpha_1}$=4 aa) with A'-A' ($\mathbf{v_2}$) vs. A-GEMS ($\boldsymbol{\alpha_1}$=4 aa) with A'-A' ($\mathbf{v_3}$) | 2.745 | ns (p=0.1424) | - | - | 8 |
| S6f | A-GEMS ($\boldsymbol{\alpha_2}$=8 aa) with A'-A' vs. A-GEMS ($\boldsymbol{\alpha_2}$=8 aa) with A'-A' ($\mathbf{v_2}$) vs. A-GEMS ($\boldsymbol{\alpha_2}$=8 aa) with A'-A' ($\mathbf{v_3}$) | 10.13 | * (p=0.0119) | A-GEMS ($\boldsymbol{\alpha_2}$=8 aa) with A'-A' vs. A-GEMS ($\boldsymbol{\alpha_2}$=8 aa) with A'-A' ($\mathbf{v_2}$) | * (p=0.0181) Tukey | 8 |
| | | | | A-GEMS ($\boldsymbol{\alpha_2}$=8 aa) with A'-A' vs. A-GEMS ($\boldsymbol{\alpha_2}$=8 aa) with A'-A' ($\mathbf{v_3}$) | * (p=0.0195) Tukey | |
| | | | | A-GEMS ($\boldsymbol{\alpha_2}$=8 aa) with A'-A' ($\mathbf{v_2}$) vs. A-GEMS ($\boldsymbol{\alpha_2}$=8 aa) with A'-A' ($\mathbf{v_3}$) | ns (p=0.9975) Tukey | |
| S6f | A-GEMS ($\boldsymbol{\alpha_3}$=27 aa) with A'-A' vs. A-GEMS ($\boldsymbol{\alpha_3}$=27 aa) with A'-A' ($\mathbf{v_2}$) vs. A-GEMS ($\boldsymbol{\alpha_3}$=27 aa) with A'-A' ($\mathbf{v_3}$) | 13.51 | ** (p=0.006) | A-GEMS ($\boldsymbol{\alpha_3}$=27 aa) with A'-A' vs. A-GEMS ($\boldsymbol{\alpha_3}$=27 aa) with A'-A' ($\mathbf{v_2}$) | ns (p=0.0519) Tukey | 8 |
| | | | | A-GEMS ($\boldsymbol{\alpha_3}$=27 aa) with A'-A' vs. | ** (p=0.005) Tukey | |

| | | | | | | |
|---|---|---|---|---|---|---|
| | | | | A-GEMS ($\alpha_3$=27 aa) with A’-A’ ($v_3$) | | |
| | | | | A-GEMS ($\alpha_3$=27 aa) with A’-A’ ($v_2$) vs. A-GEMS ($\alpha_3$=27 aa) with A’-A’ ($v_3$) | ns (p=0.1632) Tukey | |
| 4a | A-GEMS ($\alpha_2$=8 aa) with A’-A’ vs. A-GEMS ($\alpha_2$=8 aa) with B’-B’ vs. A-GEMS ($\alpha_2$=8 aa) with C’-C’ | 285.1 | *** (p<0.0001) | A-GEMS ($\alpha_2$=8 aa) with A’-A’ vs. A-GEMS ($\alpha_2$=8 aa) with B’-B’ | *** (p<0.0001) Tukey | 8 |
| | | | | A-GEMS ($\alpha_2$=8 aa) with A’-A’ vs. A-GEMS ($\alpha_2$=8 aa) with C’-C’ | *** (p<0.0001) Tukey | |
| | | | | A-GEMS ($\alpha_2$=8 aa) with B’-B’ vs. A-GEMS ($\alpha_2$=8 aa) with C’-C’ | ns (p=0.9994) Tukey | |
| 4a | B-GEMS ($\alpha_2$=8 aa) with A’-A’ vs. B-GEMS ($\alpha_2$=8 aa) with B’-B’ vs. B-GEMS ($\alpha_2$=8 aa) with C’-C’ | 1155 | *** (p<0.0001) | B-GEMS ($\alpha_2$=8 aa) with A’-A’ vs. B-GEMS ($\alpha_2$=8 aa) with B’-B’ | *** (p<0.0001) Tukey | 8 |
| | | | | B-GEMS ($\alpha_2$=8 aa) with A’-A’ vs. B-GEMS ($\alpha_2$=8 aa) with C’-C’ | ns (p=0.7867) Tukey | |
| | | | | B-GEMS ($\alpha_2$=8 aa) with B’-B’ vs. B-GEMS ($\alpha_2$=8 aa) with C’-C’ | *** (p<0.0001) Tukey | |
| 4a | C-GEMS ($\alpha_2$=8 aa) with A’-A’ vs. C-GEMS ($\alpha_2$=8 aa) with B’-B’ vs. C-GEMS ($\alpha_2$=8 aa) with C’-C’ | 5257 | *** (p<0.0001) | C-GEMS ($\alpha_2$=8 aa) with A’-A’ vs. C-GEMS ($\alpha_2$=8 aa) with B’-B’ | ns (p=0.8865) Tukey | 8 |
| | | | | C-GEMS ($\alpha_2$=8 aa) with A’-A’ vs. C-GEMS ($\alpha_2$=8 aa) with C’-C’ | *** (p<0.0001) Tukey | |
| | | | | C-GEMS ($\alpha_2$=8 aa) with B’-B’ vs. C-GEMS ($\alpha_2$=8 aa) with C’-C’ | *** (p<0.0001) Tukey | |
| S10 | B-GEMS ($\alpha_2$=8 aa) with A’-A’ vs. A-GEMS ($\alpha_2$=8 aa) with A’-A’ vs. | 696.2 | *** (p<0.0001) | A-GEMS ($\alpha_2$=8 aa) with A’-A’ vs. | ns (p=0.3039) Bonferroni | 8 |

| | | | | | | |
|---|---|---|---|---|---|---|
| | A-GEMS ($\boldsymbol{\alpha}_2$=8 aa) with<br>A’-A’ and B’-B’ and C’-C | | | A-GEMS ($\boldsymbol{\alpha}_2$=8 aa)<br>with A’-A’ and B’-B’<br>and C’-C | | |
| S14b | A-GEMS ($\boldsymbol{\alpha}_2$=8 aa) with no ligand<br>vs.<br>A-GEMS ($\boldsymbol{\alpha}_2$=8 aa) with A’-Γ<br>vs.<br>A-GEMS ($\boldsymbol{\alpha}_2$=8 aa) with A’-Γ’ | 10.8 | *<br>(p=0.0103) | A-GEMS ($\boldsymbol{\alpha}_2$=8 aa)<br>with no ligand<br>vs.<br>A-GEMS ($\boldsymbol{\alpha}_2$=8 aa)<br>with A’-Γ | ns (p>0.05) | 8 |
| | | | | A-GEMS ($\boldsymbol{\alpha}_2$=8 aa)<br>with no ligand<br>vs.<br>A-GEMS ($\boldsymbol{\alpha}_2$=8 aa)<br>with A’-Γ’ | ** (p<0.01)<br>Dunnett | |
| S17e | A-GEMS: B-GEMS ($\boldsymbol{\alpha}_2$=8 aa)<br>vs.<br>A-GEMS ($\boldsymbol{\alpha}_2$=8 aa) with A’-A’<br>(bacterial)<br>vs.<br>A-GEMS ($\boldsymbol{\alpha}_2$=8 aa) with A’-A’<br>(mammalian)<br>vs.<br>A-GEMS ($\boldsymbol{\alpha}_2$=8 aa) with A’<br>vs.<br>A-GEMS ($\boldsymbol{\alpha}_2$=8 aa) | 2140 | ***<br>(p<0.0001) | A-GEMS ($\boldsymbol{\alpha}_2$=8 aa)<br>with A’-A’ (bacterial)<br>vs.<br>A-GEMS ($\boldsymbol{\alpha}_2$=8 aa)<br>with A’-A’<br>(mammalian) | *** (p<0.0001)<br>Bonferroni | 14 |
| 5b | receivers with ligand<br>vs.<br>receivers with senders<br>(+ dox)<br>vs.<br>receivers with senders<br>(- dox)<br>vs.<br>receivers with HEK293S GnTi$^-$<br>TetR<br>(+ dox) | 230.8 | ***<br>(p<0.0001) | receivers with ligand<br>vs.<br>receivers with<br>senders<br>(+ dox) | *** (p<0.0001)<br>Tukey | 11 |
| | | | | receivers with ligand<br>vs.<br>receivers with<br>senders<br>(- dox) | *** (p<0.0001)<br>Tukey | |
| | | | | receivers with ligand<br>vs.<br>receivers with<br>HEK293S GnTi$^-$ TetR<br>(+ dox) | *** (p<0.0001)<br>Tukey | |
| | | | | receivers with<br>senders<br>(+ dox)<br>vs.<br>receivers with<br>senders<br>(- dox) | *** (p=0.0002)<br>Tukey | |
| | | | | receivers with<br>senders<br>(+ dox)<br>vs.<br>receivers with<br>HEK293S GnTi$^-$ TetR<br>(+ dox) | *** (p<0.0001)<br>Tukey | |
| | | | | receivers with<br>senders<br>(- dox) | ns (p=0.0569)<br>Tukey | |

| | | | | | | |
|---|---|---|---|---|---|---|
| | | | | vs.<br>receivers with HEK293S GnTi$^{-}$ TetR (+ dox) | | |
| 5d | receivers with senders 1 & senders 2<br>vs.<br>receivers with senders 1 & control<br>vs.<br>receivers with senders 2 & control<br>vs.<br>receivers with control | 185.4 | ***<br>(p<0.0001) | receivers with senders 1 & senders 2<br>vs.<br>receivers with senders 1 & control | *** (p<0.0001)<br>Tukey | 11 |
| | | | | receivers with senders 1 & senders 2<br>vs.<br>receivers with senders 2 & control | *** (p<0.0001)<br>Tukey | |
| | | | | receivers with senders 1 & senders 2<br>vs.<br>receivers with control | *** (p<0.0001)<br>Tukey | |
| | | | | receivers with senders 1 & control<br>vs.<br>receivers with senders 2 & control | * (p<0.05)<br>Tukey | |
| | | | | receivers with senders 1 & control<br>vs.<br>receivers with control | *** (p<0.0001)<br>Tukey | |
| | | | | receivers with senders 2 & control<br>vs.<br>receivers with control | *** (p<0.0001)<br>Tukey | |
| S25b | receivers with A’-A’ ligand<br>vs.<br>receivers with senders 1 & senders 2<br>vs.<br>receivers with control | 243.4 | ***<br>(p<0.0001) | receivers with A’-A’ ligand<br>vs.<br>receivers with senders 1 & senders 2 | ns (p>0.05)<br>Tukey | 8 |
| | | | | receivers with A’-A’ ligand<br>vs.<br>receivers with control | *** (p<0.0001)<br>Tukey | |
| | | | | receivers with senders 1 & senders 2<br>vs.<br>receivers with control | *** (p<0.0001)<br>Tukey | |

**Supplementary Table S7: DNA and protein sequences for linker $l_x$ spanning two CCs for expression of dipeptide ligand.** The four different linkers ($l_x$) used in this work to span two CC modules (see Supplementary Fig. S5b). For linker-length optimization (Fig. 3), linkers $l_1$, $l_2$, $l_3$[4,5] were utilized. For ligands A'Γ and A'Γ' used to construct AND gate logic, linker $l_4$ was used to span the two unique CC modules (Fig. 4b,d).

| ID | DNA Sequence (5' to 3') | Amino acid sequence (N-term to C-term) |
|---|---|---|
| $l_1$ | AGTGGATCCTCGGGATCATCAGGATCCTCTGGGAGCTCA GGATCCTCGGGGAGCTCTGGATCCAGCGGTTCAAGCGGA TCCTCAGGGAGCTCTGGATCCAGTGGTAGTTCGGGATCC AGCGGATCTAGTGGATCCTCTGGTAGTTCGGGATCCAGT GGTAGCCCGCGGATGGTGAGCAAGGGC | SGSSGSSGSSGSSGSSGSSGSSGSSGSSGSSGSS GSSGSSGSSGSSGSPRMVSKG |
| $l_2$ | TGGGAATTTCATCACGGGGGTTCGGGTGGTTCAGGCGGC TCAGGAGGCTCCGGGGGTTCCGGAGGGAGCGGTGCTGA AGCCGCAGCCAAGGAAGCAGCAGCTAAAGAGGCCGCTG CGAAGGAAGCTGCCGCAAAGGAGGCGGCGGCGAAAGA GGCGGCAGCAAAAGCCGGATCTGGTGGCAGTGGTGGCT CCGGCGGGTCAGGTGGCAGCGGGGGATCAGGAG | WEFHHGGSGGSGGSGGSGGSGGSGAEAAAKEAA AKEAAAKEAAAKEAAAKEAAAKAGSGGSGGSGGS GGSGGSG |
| $l_3$ | TGGGAATTTCATCACGGGGGTTCGGGTGGTTCAGGCGGC TCAGGAGGCTCCGGGGGTTCCGGAGGGAGCGGTGCTGA AGCCGCAGCCAAGGAAGCAGCAGCTAAAGAGGCCGCTG CGAAGGAAGCTGCCGCAAAGGAGGCGGCGGCGAAAGA GGCGGCAGCAAAAGCCGGATCTGGTGGCAGTGGTGGCT CCGGTGGGTCAGGTGGCAGCGGGGGATCAGGAGCTGAG GCAGCCGCCAAAGAGGCTGCGGCCAAGGAGGCCGCCGC TAAAGAAGCCGCGGCAAAAGAGGCAGCGGCAAAGGAAG CGGCTGCGAAAGCCGGAAGTGGTGGGTCGGGCGGCTCC GGTGGCTCAGGCGGTCTGTGG | WEFHHGGSGGSGGSGGSGGSGGSGAEAAAKEAA AKEAAAKEAAAKEAAAKEAAAKAGSGGSGGSGGS GGSGGSGAEAAAKEAAAKEAAAKEAAAKEAAAKEA AAKAGSGGSGGSGGSGGLW |
| $l_4$ | TGGGAATTTCATCACGGGGGTTCGGGTGGTTCAGGCGGC | WEFHHGGSGGSGG |

**Supplementary Table S8: DNA sequence of STAT3 recognition element for the engineering of the reporter gene.** Overview of nucleotide sequence of signal transducer and activator of transcription 3 (STAT3) recognition element with four STAT3 binding sites (STAT3 BS) and a minimal cytomegalovirus promoter (minCMV). STAT3 BS in blue; minCMV in red.

| DNA Sequence (5' to 3') |
|---|
| GGTTCCCGTAAATGCATCAGGTTCCCGTAAATGCATCAGGTTCCCGTAAATGCATCAGGTTCCCGTAAATGCATCACGGGTC GAGGTAGGCGTGTACGGTGGGAGGCCTATATAAGCAGAGCTCGTTTAGTGAACCGTCAGATC |

**Supplementary Table S9: Engineered plasmids used in the present study.** Overview of engineered plasmids, including plasmid backbone and gene insert.

| Plasmid ID | Plasmid Backbone | Gene insert | Figure-reference |
|---|---|---|---|
| pET28a(+)-SUMO-A' | pET28a(+)-$P_{T7}$ | SUMO-A' CC | Figure 2, Supplementary Fig. S4 |
| pET28a(+)-SUMO-A'($v_1$) | pET28a(+)-$P_{T7}$ | SUMO-A'.C4G CC ($v_1$) | Supplementary Fig. S6 |
| pET28a(+)-SUMO-A'($v_2$) | pET28a(+)-$P_{T7}$ | SUMO-A'.S7P CC ($v_2$) | Supplementary Fig. S6 |
| pET28a(+)-SUMO-A'($v_3$) | pET28a(+)-$P_{T7}$ | SUMO-A'.S7P.Link CC ($v_3$) | Supplementary Fig. S6 |
| pET28a(+)-SUMO-A'-$l_1$-A' | pET28a(+)-$P_{T7}$ | SUMO-A' CC-$l_1$-A' CC | Figure 3, Supplementary Fig. S7 |
| pET28a(+)-SUMO-A'-$l_2$-A' | pET28a(+)-$P_{T7}$ | SUMO-A' CC-$l_2$-A' CC | Figure 3, Figure 4, Supplementary Fig. S7, Supplementary Fig. S10 |
| pET28a(+)-SUMO-A'-$l_3$-A' | pET28a(+)-$P_{T7}$ | SUMO-A' CC-$l_3$-A' CC | Figure 3, Supplementary Fig. S7 |

| | | | |
|---|---|---|---|
| pET28a(+)-SUMO-B’-$l_2$-B’ | pET28a(+)-$P_{T7}$ | SUMO-B’ CC-$l_2$-B’ CC | Figure 4, Supplementary Fig. S10 |
| pET28a(+)-SUMO-Γ’-$l_2$-Γ’ | pET28a(+)-$P_{T7}$ | SUMO-Γ’ CC-$l_2$-Γ’ CC | Figure 4, Supplementary Fig. S10 |
| pET28a(+)-SUMO-A’-$l_2$-B’ | pET28a(+)-$P_{T7}$ | SUMO-A’ CC-$l_2$-B’ CC | Figure 4, Supplementary Fig. S11 |
| pET28a(+)-SUMO-A’-$l_2$-Γ’ | pET28a(+)-$P_{T7}$ | SUMO-A’ CC-$l_2$-Γ’ CC | Figure 4, Supplementary Fig. S11 |
| pET28a(+)-SUMO-A’-$l_4$-Γ | pET28a(+)-$P_{T7}$ | SUMO-A’ CC-$l_4$-Γ CC | Figure 4, Supplementary Fig. S15 |
| pET28a(+)-SUMO-A’-$l_4$-Γ’ | pET28a(+)-$P_{T7}$ | SUMO-A’ CC-$l_4$-Γ’ CC | Figure 4, Supplementary Fig. S15 |
| pLeo-A-GEMS | pLeo619-$P_{SV40}$ | A CC-GEMS | Figure 1, Figure 2, Supplementary Fig. S3, Supplementary Fig. S6 |
| pLeo-A-$\alpha_1$-GEMS | pLeo619-$P_{SV40}$ | A CC-$\alpha_1$ -GEMS | Figure 1, Figure 2, Supplementary Fig. S3, Supplementary Fig. S6 |
| pLeo-A-$\alpha_2$-GEMS | pLeo619-$P_{SV40}$ | A CC-$\alpha_2$ -GEMS | Figure 1, Figure 2, Figure 3, Figure 4, Supplementary Fig. S3, Supplementary Fig. S6, Supplementary Fig. S10, Supplementary Fig. S11, Supplementary Fig. S15, Supplementary Fig. S17, Supplementary Fig. S20 |
| pLeo-A-$\alpha_3$-GEMS | pLeo619-$P_{SV40}$ | A CC-$\alpha_3$-GEMS | Figure 1, Figure 2, Supplementary Fig. S3, Supplementary Fig. S6 |
| pLeo-A’-GEMS | pLeo619-$P_{SV40}$ | A’-CC-GEMS | Figure 1, Supplementary Fig. S3 |
| pLeo-A’-$\alpha_1$-GEMS | pLeo619-$P_{SV40}$ | A’ CC-$\alpha_1$-GEMS | Figure 1, Supplementary Fig. S3 |
| pLeo-A’-$\alpha_2$-GEMS | pLeo619-$P_{SV40}$ | A’ CC-$\alpha_2$-GEMS | Figure 1, Supplementary Fig. S3, Supplementary Fig. S20 |
| pLeo-A’-$\alpha_3$-GEMS | pLeo619-$P_{SV40}$ | A’ CC-$\alpha_3$-GEMS | Figure 1, Supplementary Fig. S3 |
| pLeo-B-$\alpha_2$-GEMS | pLeo619-$P_{SV40}$ | B CC-$\alpha_2$-GEMS | Figure 1, Figure 2, Figure 4, Supplementary Fig. S3, Supplementary Fig. S6, Supplementary Fig. S10, Supplementary Fig. S11 |
| pLeo-Γ-$\alpha_2$-GEMS | pLeo619-$P_{SV40}$ | Γ CC-$\alpha_2$-GEMS | Figure 4, Supplementary Fig. S11 |
| pHR-EGFPligand | pHR-$P_{SFFV}$ | EGFP-PDGFR | Figure 1, Figure2, Supplementary Fig. S3 (pHR_EGFPligand was a gift from Wendell Lim (Addgene plasmid # 79129 ; http://n2t.net/addgene:79129 ; RRID:Addgene_79129))[6] |
| pHR-SUMO-A’-$l_2$-A’-IRES-EmGFP | pHR-$P_{CMV-TetO2}$ | SUMO-A’ CC-$l_2$-A’ CC-IRES-EmGFP | Figure 5, Supplementary Fig. S17, Supplementary Fig. S18 |

| | | | |
|---|---|---|---|
| pHR- SEAP-PGK-mCherry | pHR-$P_{CMV}$ | STAT3 recognition element-SEAP-PGK-mCherry | Figure 5, Supplementary Fig. S20, Supplementary Fig. S21, Supplementary Fig. S25, Supplementary Fig. S26 |
| pHR-A-$\alpha_2$-GEMS | pHR-$P_{CMV}$ | A CC-$\alpha_2$-GEMS | Figure 5, Supplementary Fig. S21, Supplementary Fig. S25, Supplementary Fig. S26 |
| pHR-SUMO-A'-$I_4$-Γ-IRES-EmGFP | pHR-$P_{CMV\text{-}TetO2}$ | SUMO-A' CC-$I_4$-Γ CC-IRES-EmGFP | Figure 5, Supplementary Fig. S23, Supplementary Fig. S25, Supplementary Fig. S26 |
| pHR-SUMO-A'-$I_4$-Γ'-IRES-EmGFP | pHR-$P_{CMV\text{-}TetO2}$ | pHR-SUMO-A' CC-$I_4$-Γ' CC-IRES-EmGFP | Figure 5, Supplementary Fig. S24, Supplementary Fig. S25, Supplementary Fig. S26 |

**Supplementary Table S10: Receiver : Sender 1 : Sender 2 ratios for co-culture experiments.** Overview of the ratios of Receiver : Sender 1 : Sender 2 cell populations, as well as cell density numbers for Receiver, Sender 1, Sender 2 and control (untransduced HEK293S GnTi$^-$) cells presented in Supplementary Figure S26. Cell numbers are reported for seeding in a 24 well plate.

| **Ratio (Receivers : Senders 1 : Senders 2)** | **Receivers** | **Senders 1** | **Senders 2** |
|---|---|---|---|
| 1:1:1 | $1.2 \times 10^5$ | $1.2 \times 10^5$ | $1.2 \times 10^5$ |
| 3:1:1 | $3.6 \times 10^5$ | $1.2 \times 10^5$ | $1.2 \times 10^5$ |
| 4:1:1 | $3.2 \times 10^5$ | $0.8 \times 10^5$ | $0.8 \times 10^5$ |
| 5:1:1 | $4.0 \times 10^5$ | $0.8 \times 10^5$ | $0.8 \times 10^5$ |
| 10:1:1 | $4.0 \times 10^5$ | $0.4 \times 10^5$ | $0.4 \times 10^5$ |

### Supplementary Figures

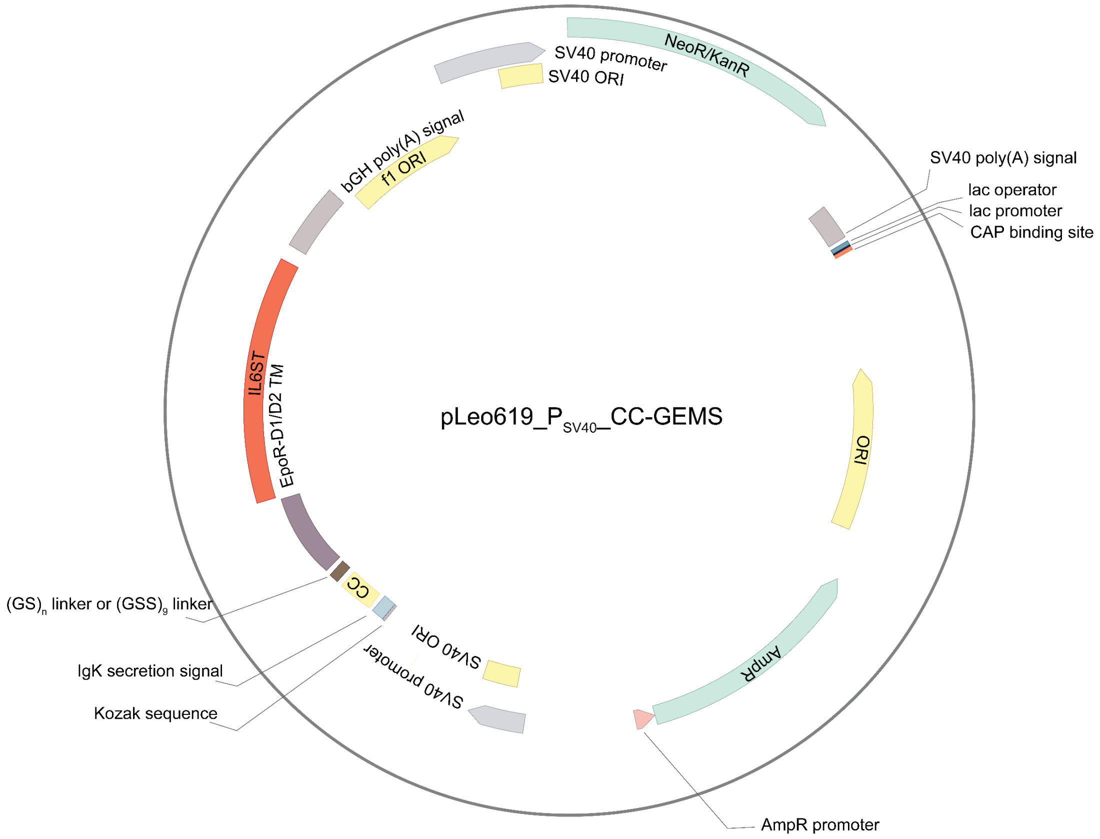


**Supplementary Fig. S1: Plasmid map of pLeo619 expression vector for transient mammalian expresion of the CC functionalized GEMS receptor.** The CC-GEMS receptor consists of a selected CC (see Supplementary Table S1), a linker composed of GS or GGS repeats for a total of zero, four, eight or 27 aa length ($(GS)_n$ linker, where n=0, 2, 4 or $(GSS)_9$ linker) and the EpoR transmembrane domain (EpoR-D1/D2 TM) fused to the intracellular signal transduction domains of IL-6RB (interleukin 6 receptor B; IL6ST), collectively referred to as the GEMS receptor. The construct is under the control of a mammalian SV40 promoter taken from the Simian vacuolating virus 40. Ampicillin resistance gene (AmpR) is available for bacterial selection downstream an AmpR promoter. ORI, origin of replication; CAP, catabolite binding protein; NeoR/KanR; neomycin/kanamycin resistance; bGH, bovine growth hormone.

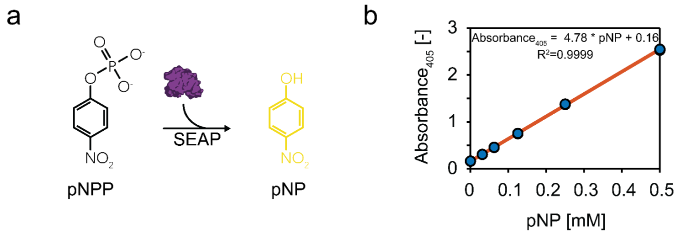


**Supplementary Fig. S2: Calibration curve to determine pNPP conversion by SEAP. a** SEAP catalyses the hydrolysis of the chromogenic substrate p-Nitrophenylphosphate (pNPP) to p-Nitrophenol (pNP), the absorbance of which can be measured at 405 nm. **b** To convert absorption units to amount of pNP, we titrated known concentrations of pNP (0 mM, 0.03125 mM, 0.0625 mM, 0.125 mM, 0.25 mM and 0.5 mM), measured absorbance units (405 nm at 25°C) and fitted the data points using a linear regression curve. $Absorbance_{405} = 4.78 * [pNP] + 0.16$, $R^2=0.9999$. Individual data points represent technical replicates (n=3).

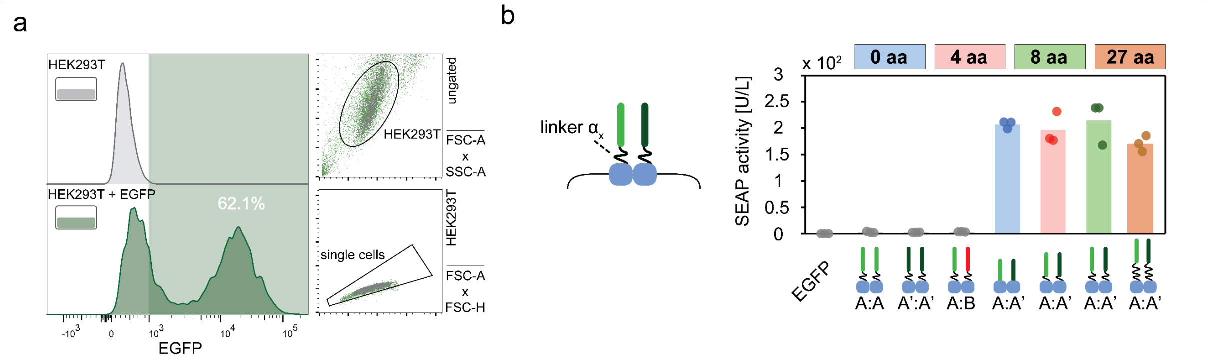


**Supplementary Fig. S3: Cognate CC-receptor heterodimerization. a** Flow cytometry analysis of EGFP transfected HEK293T cells (green; see Methods) and control (untransduced HEK293T; grey), showing the gating strategy on the right (dot plots; FSC-A x SSC-A and FSC-A x FSC-H). From the transfected population, 62.1% of HEK293T expressed EGFP. **b** SEAP activity (in U/L) in HEK293T cells transiently transfected with receptor pairs with varied linker lengths **α$_x$** of zero, four, eight and 27 amino acids (aa; GS or GGS repeats, see Methods, Figure 1d and Supplementary Table S3). SEAP activity was incubated 48 hours following transfection (see Methods). Only the correct combination of cognate CCs (A:A') result in receptor activation, while non-cognate pairs (A:A, A':A' and A:B) result in no activation. Bars indicate mean activity; individual data points represent independent triplicates.

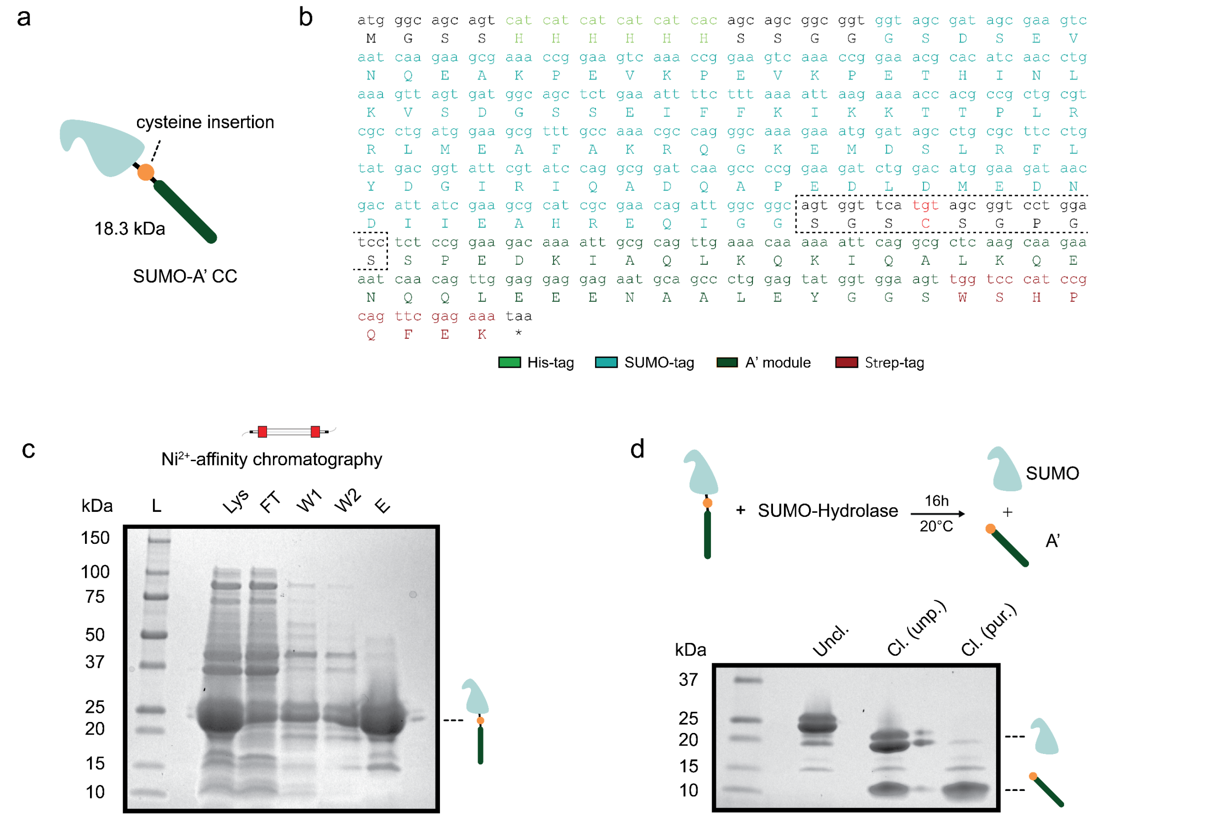


**Supplementary Fig. S4: Expression of monomeric CC peptide A'. a** Schematic representation of SUMO-A' CC fusion protein (expected size: 18.3 kDa). A cleavable SUMO-tag is N-terminally fused to monomeric CC peptide A', bearing a single cysteine. The SUMO tag carries a His-tag in the N-terminus and the A' CC carries a Strep-tag in the C-terminus (see Methods and Figure 2b). **b** DNA and protein sequence of the SUMO-A' CC fusion protein. The single-letter amino acid code is shown in uppercase below the corresponding DNA sequence. The His-tag is shown in light green, the SUMO-tag in light blue, the cysteine in red, the A' CC module in dark green and the Strep-tag in red. **c** SUMO-A' CC fusion protein was expressed in E. coli (see Methods, Supplementary Table 2 and supplementary Figure S5a) and purification with $Ni^{2+}$-affinity chromatography afforded pure SUMO-A' CC fusion protein, as shown by SDS-PAGE analysis. L: molecular weight ladder, Lys: cell lysate, FT: flow through, W1: wash fraction 1, W2: wash fraction 2, E: elution fraction. **d** SUMO-A' CC fusion protein is incubated with SUMO-hydrolase for 16 h, at 20°C (upper panel), resulting in the cleavage of the SUMO tag from the A' CC (lower panel, Cl. (unp.)). Following purification with $Ni^{2+}$ affinity chromatography, CC peptide A' was recovered (Cl. (pur.)).

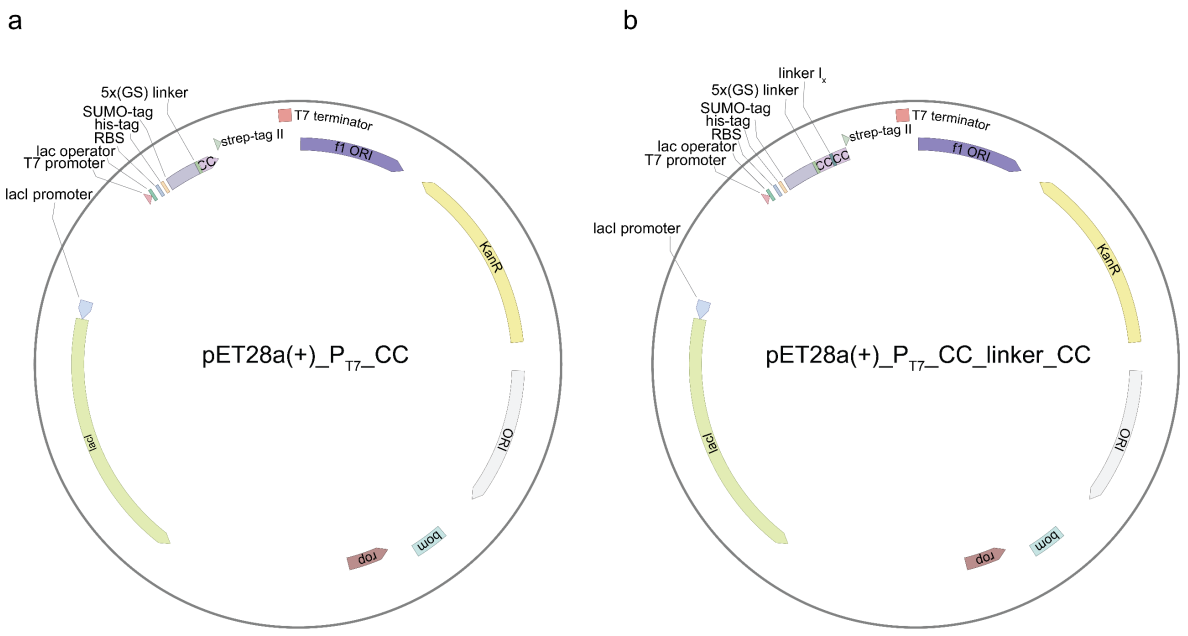


**Supplementary Fig. S5: Plasmid maps of pET28a(+) expression vector for expression of the monomeric and dimeric CC ligands. a** The selected CC is SUMO-tagged and expressed downstream of a T7 promoter. An N-terminus His-tag and a C-terminus strep-tag are incorporated for subsequent purification. **b** Two unique CC modules are spanned by a linker **$l_x$** (see Supplementary Table S7). A His-tag is engineered on the N-terminus of the SUMO-tag and a strep tag on the C-terminus of the CC-linker-CC fusing protein. A kanamycin resistance gene (KanR) is available for bacterial selection. ORI, origin of replication; bom; basis of mobility; rop, repressor of primer; RBS, ribosome binding site.

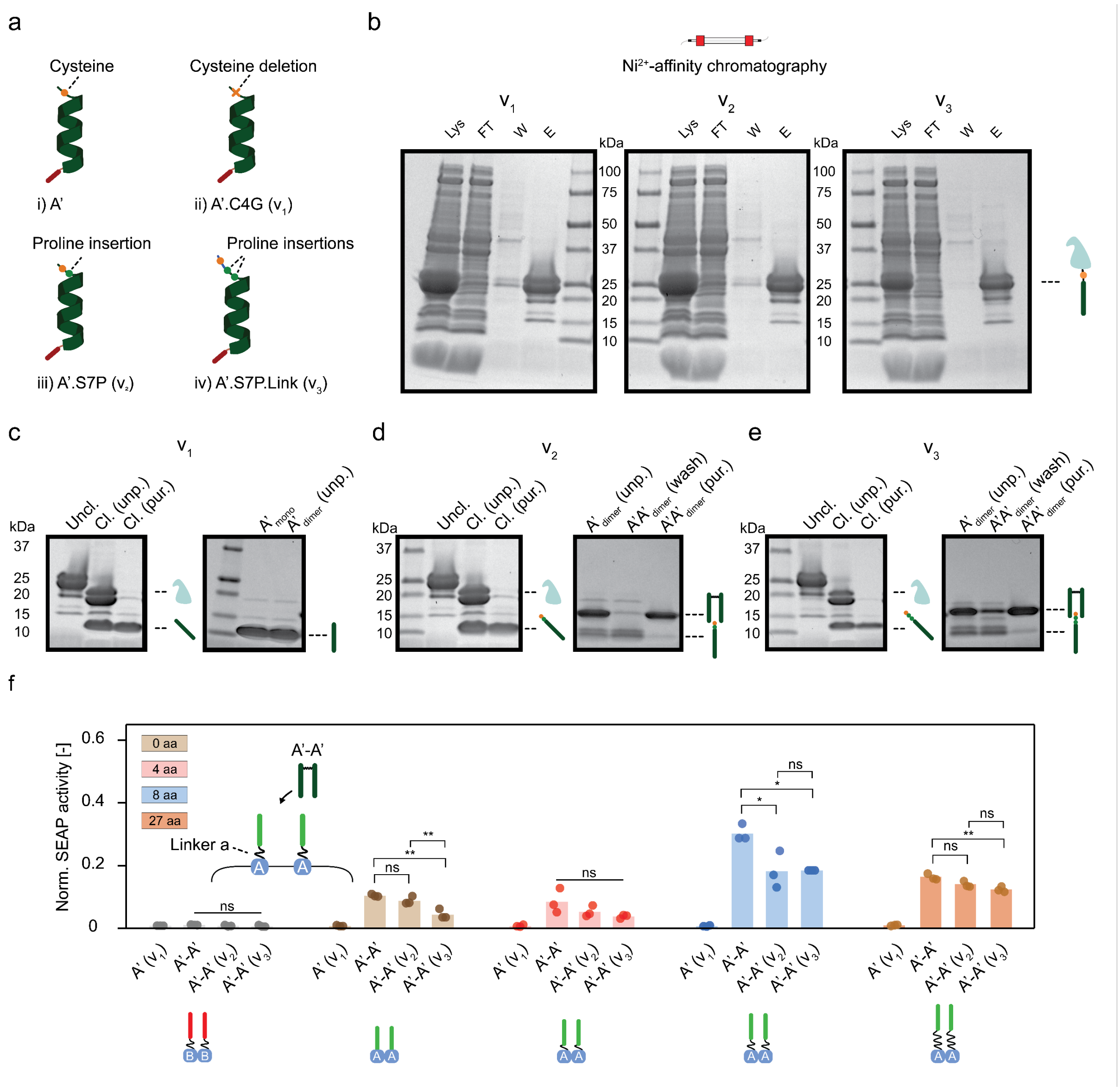


**Supplementary Fig. S6: Mutant A' CC expression, purification, and subsequent effects on receptor activation.** **a** Schematic representation of strategy to express three variants (**$v_x$**) of mutant monomeric A' CCs; i) A': original A' monomer (see Supplementary Figure S4b), ii) A'.C4G (**$v_1$**): cysteine deletion, iii) A'.S7P (**$v_2$**): mutant with the addition of a proline, iv) A'.S7P.Link (**$v_3$**): variant with addition of a small linker (see Supplementary Table S4). **b** SDS-Page analysis of the expression of the variants **$v_1$**, **$v_2$** and **$v_3$** following purification with $Ni^{2+}$-affinity chromatography. Lys: cell lysate, FT: flow through, W: wash fraction, E: elution fraction. **c** SDS-PAGE of **$v_1$** prior to SUMO-cleavage (Uncl.), after (Cl.(Unp.)) and following purification with $Ni^{2+}$-affinity chromatography (Cl.(pur.)) (left). SDS-PAGE analysis, showing peptide, following incubation with homobifunctional bismaleimide linker (right; see Methods and Figure 2c). **d** and **e** SDS-PAGE analysis of the engineering of **$v_2$** and **$v_3$** dipeptides as in **c**. Ditopic A'A' ligands were formed only when N-terminal cysteine was present. Anion exchange chromatography is able to remove unreactive A' monomers (A'A'$_{dimer}$(wash)) and recover only ditopic A'A' ligands (A'A'$_{dimer}$(pur.)). **f** Normalized SEAP activity in HEK293T cells transiently transfected with receptor homodimers (A-A receptor in light green; or B-B receptor in red) with varied linkers **$\alpha_x$** of zero, four, eight and 27 aa (GS or GGS repeats). Cells were incubated with either 0.12 µM A'-A' dipeptide mutants (A'-A', A'-A' (**$v_2$**) or A'-A' (**$v_3$**)) or 0.24 µM of monomeric A'(v1) CC for 48h and SEAP expression was measured to assess receptor activation (see Methods). Data shown are normalized on a heterodimeric A-A' receptor positive control (not shown). All mutant ditopic ligands results in reporter expression for cells expressing a receptor bearing a compatible CC (A) for all linkers **$\alpha_x$**. Linker **$\alpha_x$** seems to play a role in expression levels of reporter because of receptor activation. Amongst mutant dipeptides, no observable difference in levels of receptor activation is seen. Monomeric CCs result in no activation. Additionally, the ditopic ligands fail to activate receptors harbouring a non-cognate CC with an 8 aa linker **$\alpha_2$** (B-B receptor in red). Bars indicate mean activity; individual data points represent independent triplicates. Normalization is done on the SEAP expression of the A:A' receptor heterodimer with linker **$\alpha_x$** of eight aa. Significance (ANOVA with Tukey multiple comparison correction) is noted above bars (see Supplementary Table S6). ns: p>0.05, *p ≤0.05, **p ≤0.01, ***p ≤0.001.

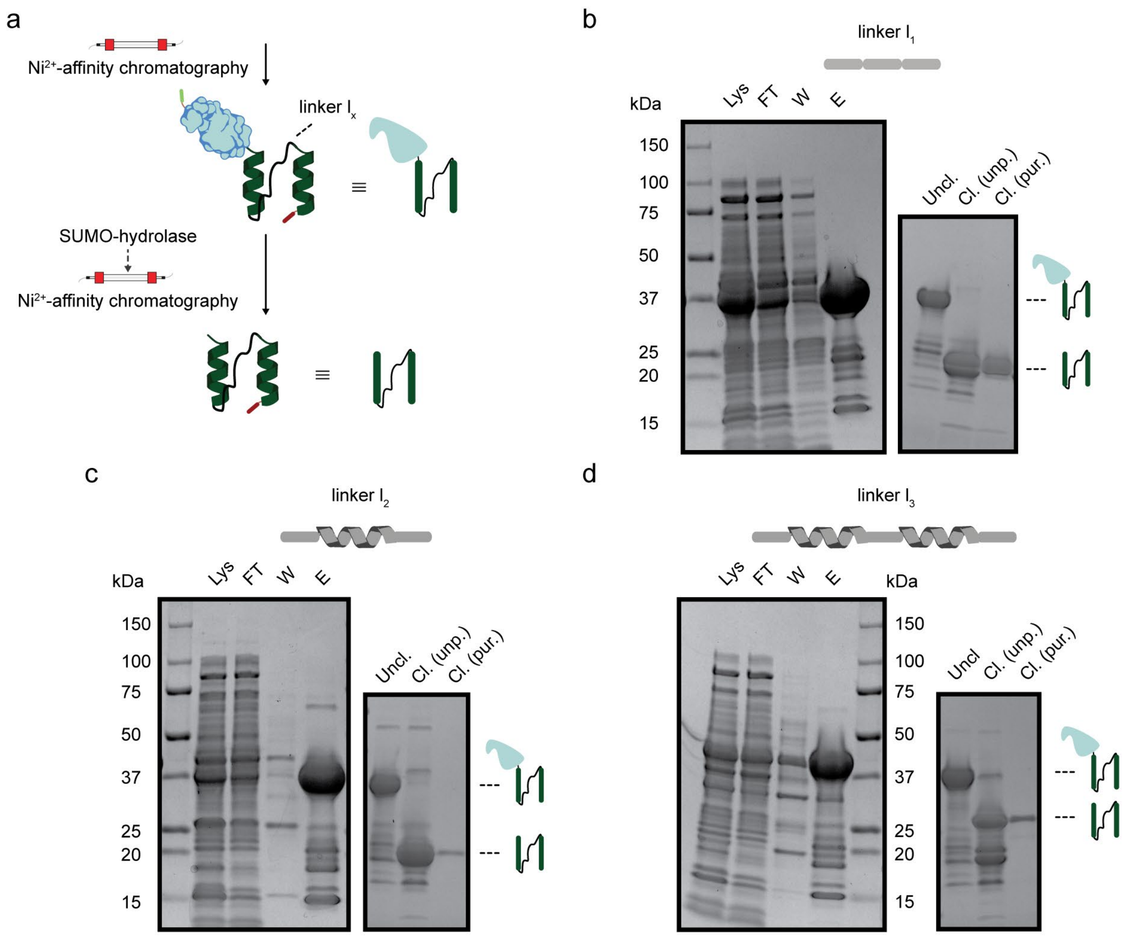


**Supplementary Fig. S7: Protein linker-based dipeptide A'-A' ligand expression and purification. a** SUMO-tagged A'-A' dipeptide carrying protein linker **l** is purified using $Ni^{2+}$-affinity chromatography. The SUMO tag is removed and A'-A' dipeptide is purified and recovered using an additional $Ni^{2+}$-affinity chromatography step (see Methods). **b, c** and **d** SDS-PAGE analysis, showing the expression of the engineered A'-A' dipeptides carrying linker **$l_1$** (**b**), **$l_2$** (**c**) and **$l_3$** (**d**), following purification with $Ni^{2+}$-affinity chromatography (left panels). Lys: cell lysate, FT: flow through, W: wash fraction, E: elution fraction. Right: Un-cleaved A'-A' CC dipeptide (Uncl.) was incubated with SUMO-hydrolase (Cl. (unp.)) to cleave of the SUMO-tag (see Methods). An additional step of $Ni^{2+}$-affinity chromatography afforded pure A'-A' CC dipeptide (Cl. (pur.)).

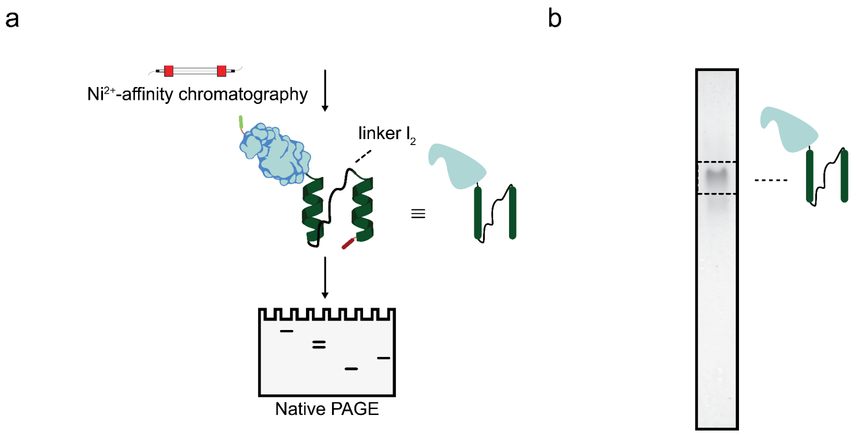


**Supplementary Fig. S8: Native PAGE of A'-A' dipeptide. a** SUMO-tagged A'-A' dipeptide with protein linker **l₂** is purified using $Ni^{2+}$-affinity chromatography and subsequently run on a Native-PAGE gel to assess ligand aggregation and oligomer formation (see Methods). **b** Native-PAGE analysis of purified A'-A' ligand showing one distinct band.

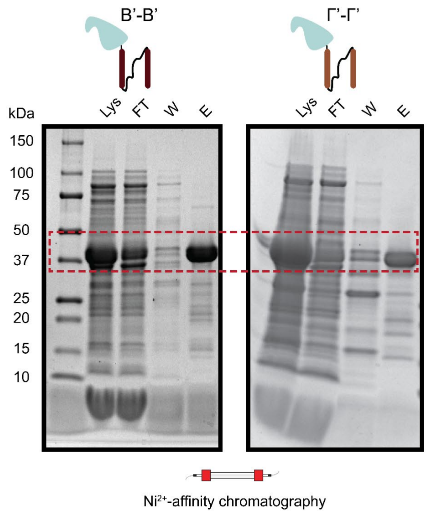


**Supplementary Fig. S9: B'-B' and Γ'-Γ' CC ligand expression and purification.** SDS-PAGE analysis, showing the expression of the engineered B'-B' (left panel) and Γ'-Γ' (right panel) CC dipeptides carrying linker **l₂**, following purification with $Ni^{2+}$-affinity chromatography. Lys: cell lysate, FT: flow through, W: wash fraction, E: elution fraction. Red inset highlights the position of the ligand on gel.

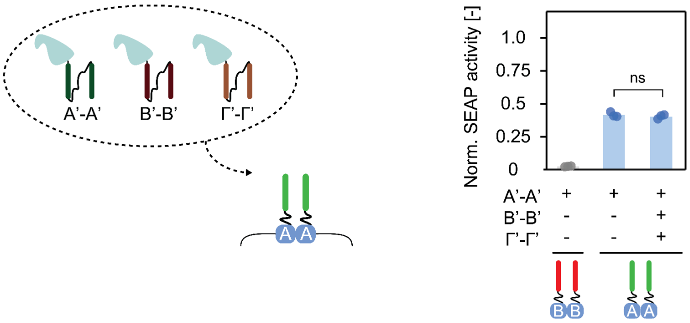


**Supplementary Fig. S10: Alternative non-cognate ligands do not inhibit activation in cells because of cognate receptor-ligand pairing.** 0.12 µM SUMO-tagged A'-A', B'-B' and Γ'-Γ' dipeptides with protein linker **l₂** were incubated with HEK293T cells transiently transfected to express A-type (green) or B-type (red) receptor with linker **α₂** (8 aa, GS repeats). Following transfection, cells were incubated with ligand for 48 hours and SEAP activity was measured (see Methods). Right panel shows the normalized SEAP activity for different receptor-ligand combination pairs. Cells expressing the A-type receptor are activated by the presence of A'-A' ligand. SEAP activity is not inhibited by the presence of B'-B' and Γ'-Γ' dipeptide ligands. Bars indicate mean activity; individual data points represent independent triplicates. Normalization is done on the SEAP expression of the A:A' receptor heterodimer with linker **α₂** of eight aa (not shown). Significance (ANOVA with Bonferroni test) is noted above bars (Supplementary Table S6). ns: p>0.05, *p ≤0.05, **p ≤0.01, ***p ≤0.001.

---

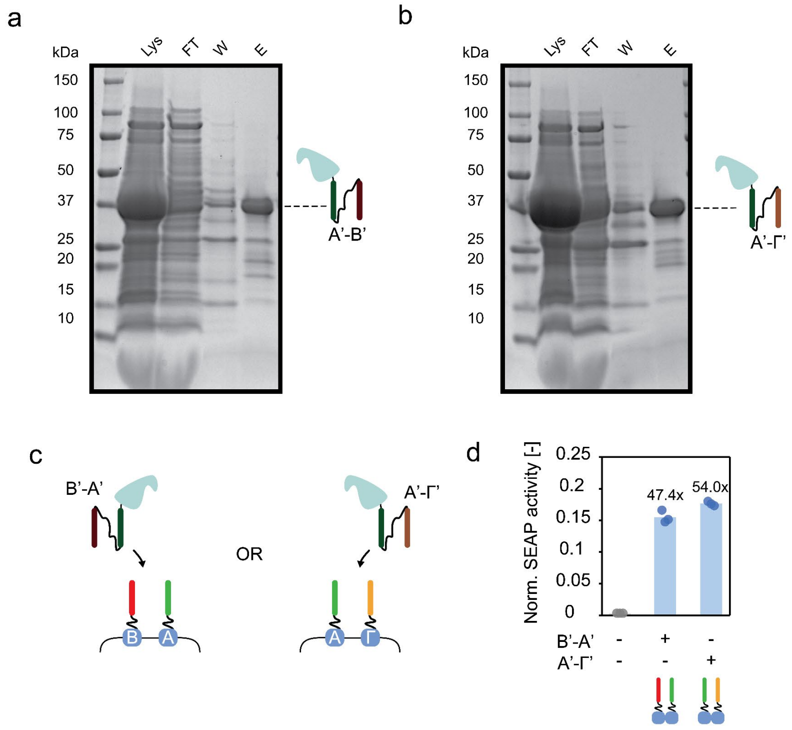


**Supplementary Fig. S11: B'-A' and A'-Γ' dipeptide ligands activate cells transfected with B-A and A-Γ receptors.** SDS-PAGE analysis, showing the expression of the engineered B'-A' (**a**) and A'-Γ' (**b**) CC dipeptides carrying linker **l₂**, following purification with $Ni^{2+}$-affinity chromatography (Methods). Lys: cell lysate, FT: flow through, W: wash fraction, E: elution fraction. **c** HEK293T cells transiently transfected with either B- and A- type or A- and Γ-type receptors, with 8 aa linker **a₂** (GS repeats) were incubated with 0.5 µM cognate SUMO-tagged dipeptide ligand B'-A' and A'-Γ' with linker **l₂** respectively for 48 hours and SEAP activity was measured (see Methods). **d** Following incubation with ligand, normalized SEAP activity was determined; which shows receptor activation for cognate ligand-receptor pairs (B'-A' ligand and B- and A-type receptor; and A'-Γ' ligand and A- and Γ-type receptors), at similar levels. The absence of ligand results in negligible activation. Bars indicate mean activity; individual data points represent independent triplicates. Normalization is done on the SEAP expression of the A:A' receptor heterodimer with linker $\alpha_2$ of eight aa (not shown). Fold change compared to cells incubated with no ligand in noted above the bars.

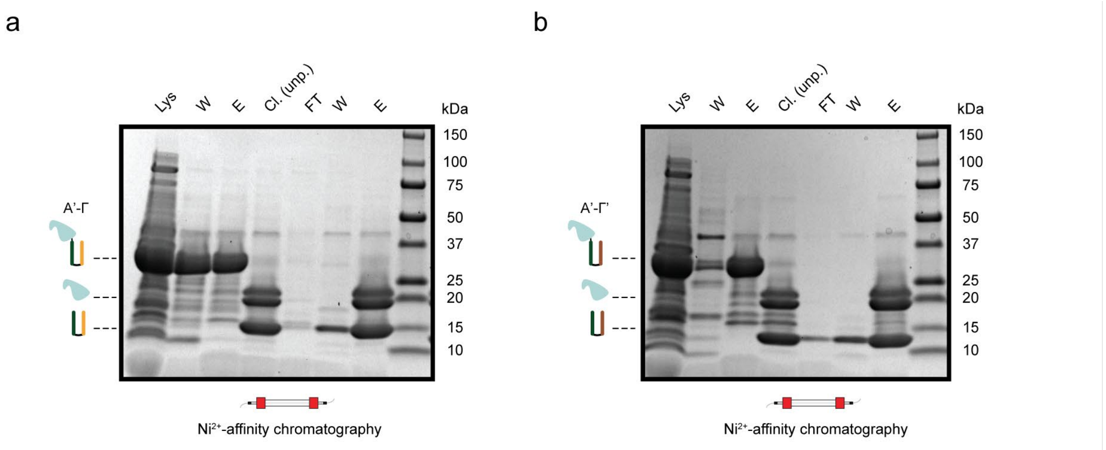


**Supplementary Fig. S12: A'-Γ and A'-Γ' dipeptide ligand expression and purification.** SDS-PAGE analysis, showing the expression of the engineered A'-Γ (**a**) and A'-Γ' (**b**) CC dipeptides carrying linker **l**$_4$, following purification with Ni$^{2+}$-affinity chromatography (Methods). Lys: cell lysate, FT: flow through, W: wash fraction, E: elution fraction. SUMO-tagged CC dipeptides were incubated with SUMO-hydrolase (Cl. (unp.)) to cleave of the SUMO-tag (see Methods).

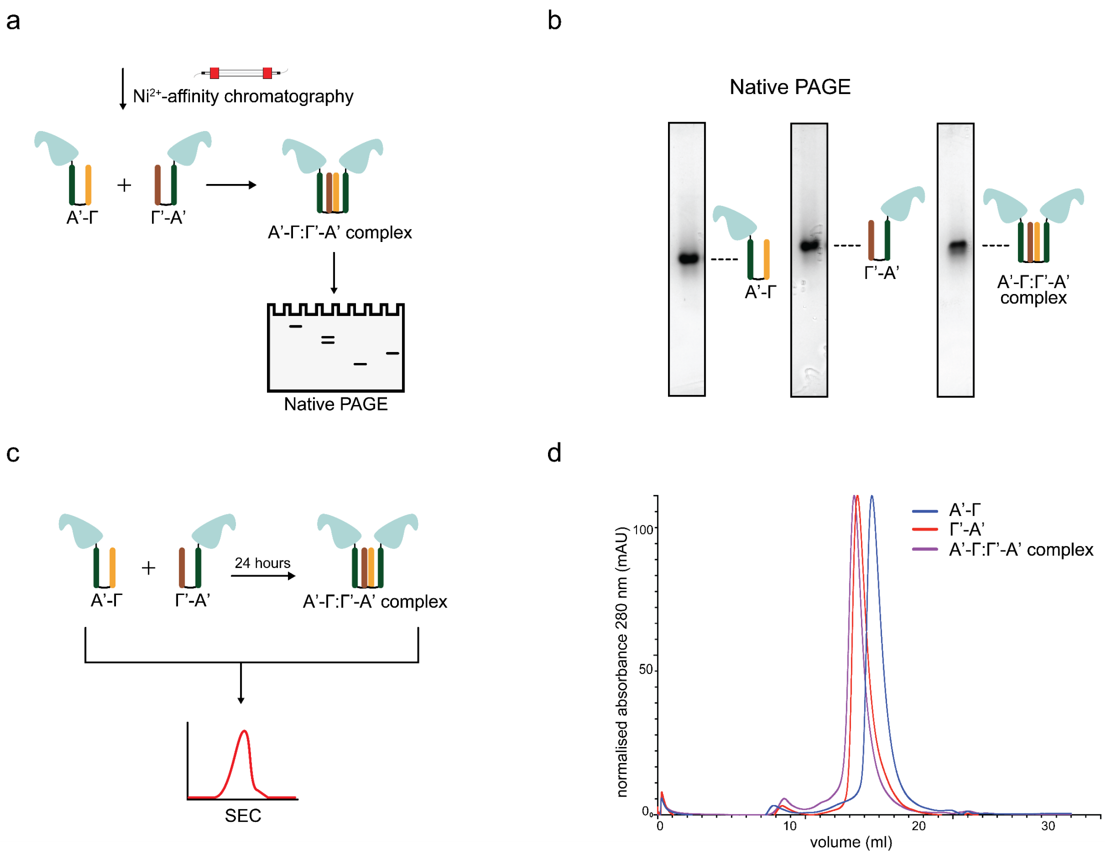

**Supplementary Fig. S13: Native PAGE of A’-Γ:Γ’-A’ complex. a** SUMO-tagged A’-Γ and A’-Γ’ dipeptides with protein linker **l₄** are purified using $Ni^{2+}$-affinity chromatography and subsequently incubated for 1 hour (see Methods). The product is run on a Native-PAGE gel to assess A’-Γ:Γ’-A’ complex formation (see Methods). **b** Native-PAGE analysis of purified A’-Γ, A’-Γ’ and A’-Γ:Γ’-A’ complex, showing a distinct band for all three products. Proteins were run in the same gel. **c** Equimolar concentrations of SUMO-tagged A’-Γ and A’-Γ’ dipeptides with protein linker **l₄** were incubated for 24 hours and were subsequently characterised using Size Exclusion Chromatography (SEC; see Methods). **d** SEC analysis for SUMO-tagged A’-Γ (blue line) and A’-Γ’ (red line) dipeptides as well as complex after incubation of one 24 hours (purple line). Incubation of dipeptides for 24 hours resulted in a distinct chromatogram peak indicating complex formation.

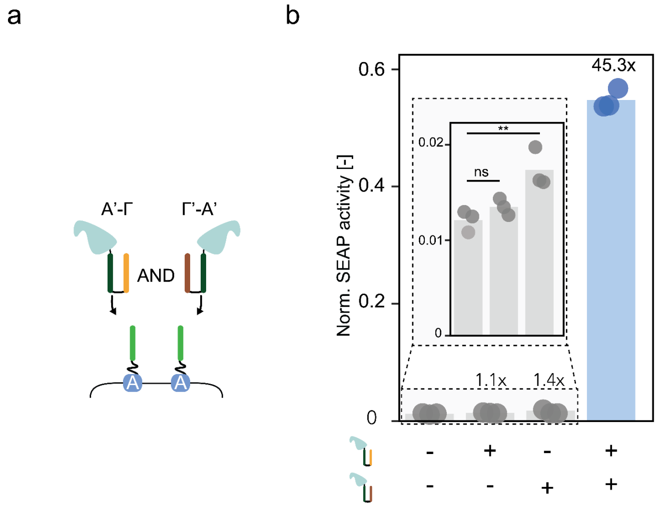


**Supplementary Fig. S14: AND gate logic based on SUMO-tagged A’-B and A’-B’ ligands, in cells expressing A-type receptor. a** HEK293T cells expressing A type receptor and reporter were incubated with either both SUMO-tagged A’-Γ and A’-Γ’ ligands or each individual A’-Γ or A’-Γ’ ligands. **b** Normalized SEAP activity in HEK293T cells transiently transfected with A-type receptor with linker **α₂** (8 aa, GS repeats), and incubated with 0.12 µM purified, SUMO-tagged ligand A’-Γ and/or Γ’-A. SEAP activity was measured after 48 hours incubation with ligand (see Methods) (see Figure 4d). Bars indicate mean activity; individual data points represent independent triplicates. Normalization is done on the SEAP expression of the A:A’ receptor heterodimer with linker **α₂** of eight aa (not shown). Inset shows the incubation with A’-Γ, Γ’-A or no ligand in a focused manner, along with a statistical analysis. Significance (ANOVA with Dunnett’s test) is noted above bars (see Supplementary Table S6). ns: $p>0.05$, *$p \leq 0.05$, **$p \leq 0.01$, ***$p \leq 0.001$. Fold change compared to cells incubated with no ligand is noted above the bars.

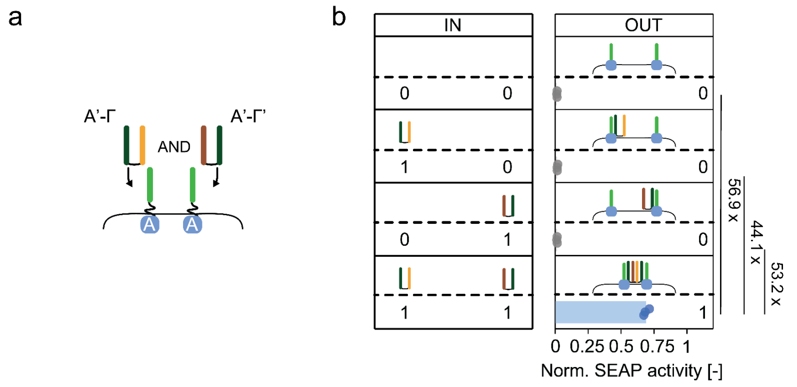


**Supplementary Fig. S15: A’-Γ and A’-Γ’ based AND gate logic in cells expressing A-type receptor. a** HEK293T cells expressing A type receptor and reporter were incubated with either both A’-Γ and A’-Γ’ ligands lacking the

SUMO tag or each individual A'-Γ or A'-Γ' ligands. **b** Normalized SEAP activity in HEK293T cells transiently transfected with receptor homodimers A-A with linker **α₂** (8 aa, GS repeats), and incubated with 0.12 µM purified ligand A'-C and/or C'-A (without the SUMO-tag). SEAP activity was measured after 48 hours incubation with ligand (see Methods). Left panel (IN; input) denotes the presence (1) or absence (0) of ditopic ligand. Right panel (OUT; output) shows normalised SEAP activity. Bars indicate mean activity; individual data points represent independent triplicates. Normalization is done on the SEAP expression of the A:A' receptor heterodimer with linker **α₂** of eight aa. Fold change is noted besides the bars.

a

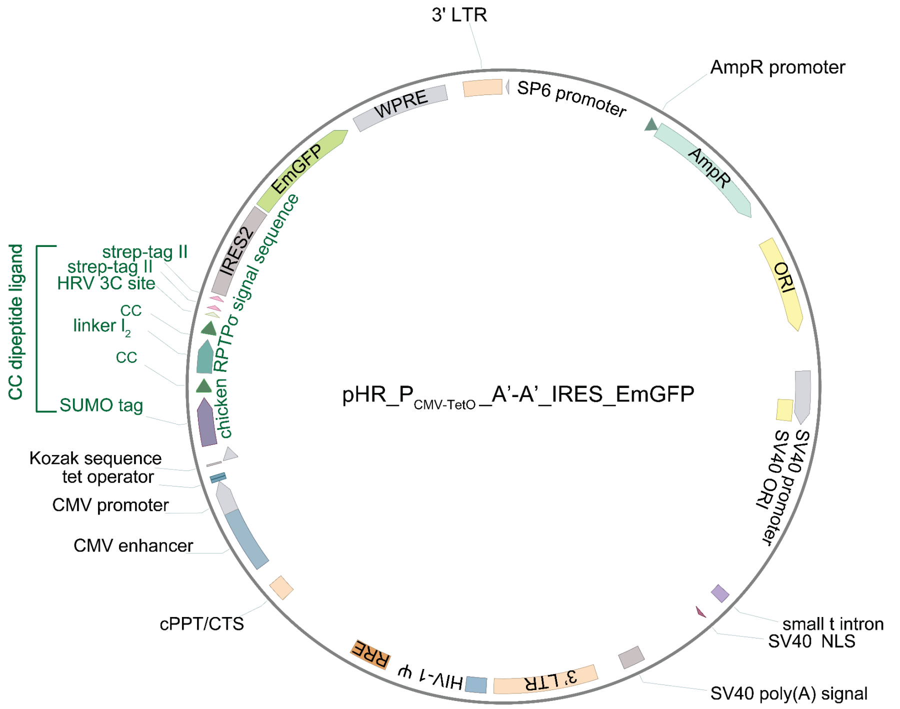


b

```
atg ggt atc ctt ccc agc cct ggg atg cct gcg ctg ctc tcc ctc gtg agc ctt ctc tcc gtg ctg ctg atg ggt tgc gta gct gaa acc ggt
 M   G   I   L   P   S   P   G   M   P   A   L   L   S   L   V   S   L   L   S   V   L   L   M   G   C   V   A   E   T   G
agc agc ggc act agt agc ggc ggt ggt agc gat agc gaa gtc aat caa gaa gcg aaa ccg gaa gtc gaa ccg gaa gtc aaa ccg gaa acg cac
 S   S   G   T   S   S   G   G   G   S   D   S   E   V   N   Q   E   A   K   P   E   V   E   P   E   V   K   P   E   T   H
atc aac ctg aaa gtt agt gat ggc agc tct gaa att ttc ttt aaa att aag aaa acc acg ccg ctg cgt cgc ctg atg gaa gcg ttt gcc aaa
 I   N   L   K   V   S   D   G   S   S   E   I   F   F   K   I   K   K   T   T   P   L   R   R   L   M   E   A   F   A   K
cgc cag ggc aaa gaa atg gat agc ctg cgc ttc ctg tat gac ggt att cgt atc cag gcg gat caa gcc ccg gaa gat ctg gac atg gaa gat
 R   Q   G   K   E   M   D   S   L   R   F   L   Y   D   G   I   R   I   Q   A   D   Q   A   P   E   D   L   D   M   E   D
aac gac att atc gaa gcg cat cgc gaa cag att ggc ggc cat atg agc ggt cct gga tcc tct ccg gaa gac aaa att gcg cag ttg aaa caa
 N   D   I   I   E   A   H   R   E   Q   I   G   G   H   M   S   G   P   G   S   S   P   E   D   K   I   A   Q   L   K   Q
aaa att cag gcg ctc aag caa gaa aat caa cag ttg gag gag gag aat gca gcc ctg gag tat ggt gga agt cga att cgg tgg gaa ttt cat
 K   I   Q   A   L   K   Q   E   N   Q   Q   L   E   E   E   N   A   A   L   E   Y   G   G   S   R   I   R   W   E   F   H
cac ggg ggt tcg ggt ggt tca ggc ggc tca gga ggc tcc ggg ggt tcc gga ggg agc ggt gct gaa gcc gca gcc aag gaa gca gca gct aaa
 H   G   G   S   G   G   S   G   G   S   G   G   S   G   G   S   G   G   S   G   A   E   A   A   A   K   E   A   A   A   K
gag gcc gct gcg aag gaa gct gcc gca aag gag gcg gcg gcg aaa gag gcg gca gca aaa gcc gga tct ggt ggc agt ggt ggc tcc ggc ggg
 E   A   A   A   K   E   A   A   A   K   E   A   A   A   K   E   A   A   A   K   A   G   S   G   G   S   G   G   S   G   G
tca ggt ggc agc ggg gga tca gga ggt acc agc ggt cct gga tcc tct ccg gaa gac aaa att gcg cag ttg aaa caa aaa att cag gcg ctc
 S   G   G   S   G   G   S   G   G   T   S   G   P   G   S   S   P   E   D   K   I   A   Q   L   K   Q   K   I   Q   A   L
aag caa gaa aat caa cag ttg gag gag gag aat gca gcc ctg gag tat ggt gga agt aag ctt tgg agc cac acg cgt ctt gag gtg ctg ttt
 K   Q   E   N   Q   Q   L   E   E   E   N   A   A   L   E   Y   G   G   S   K   L   W   S   H   T   R   L   E   V   L   F
cag gga cca gga ggt agt gga tct gct tgg agc cat cca cag ttc gaa aaa ggt gga ggt tct ggc ggt gga tca ggt gga agt gca tgg tct
 Q   G   P   G   G   S   G   S   A   W   S   H   P   Q   F   E   K   G   G   G   S   G   G   G   S   G   G   S   A   W   S
cac cct cag ttt gag aaa taa
 H   P   Q   F   E   K   *
```

chicken RPTPσ signal sequence | SUMO-tag | A' CC | linker l2 | strep-tag II

**Supplementary Fig. S16: Plasmid map and sequence of pHR expression vector for the engineering of sender cells expressing A'-A' bifunctional ligand. a** Plasmid map of lentiviral vector containing the A'-A' bifunctional ligand (denoted here in green, see Supplementary Table S3). The ligand consists of a chicken RPTPσ signal sequence for secretion, a SUMO-tag, two A' CCs spanned by a linker $\mathbf{l}_2$ (see Supplementary Table S7) and a Strep-tag II for subsequent purification (see Methods). The construct is under the control of a mammalian human cytomegalovirus (CMV) enhancer and promoter and two *TetO* operator sequences[3]. The construct is followed by an internal ribosome entry site (IRES) from encephalomyocarditis virus (EMCV) followed by an emerald green fluorescent protein (EmGFP); allowing bicistronic expression of the A'-A' dipeptide and EmGFP. WPRE: Woodchuck Hepatitis Virus posttranscriptional regulatory element, LTR: long terminal repeat, AmpR: ampicillin resistance, ORI: origin of replication, SV40: Simian vacuolating virus 40, HIV-1 ψ: retroviral psi packaging element, RRE: Rev response element, cPPT/CTS: central polypurine tract/central termination sequence. **b** DNA and protein sequence of the A'-A' bifunctional ligand. The single-letter amino acid code is shown in uppercase below the corresponding DNA sequence. Chicken RPTPσ signal sequence is shown here in red; SUMO-tag in purple; A' CC in dark green; linker $\mathbf{l}_2$ in light green; Strep-tag II in pink.

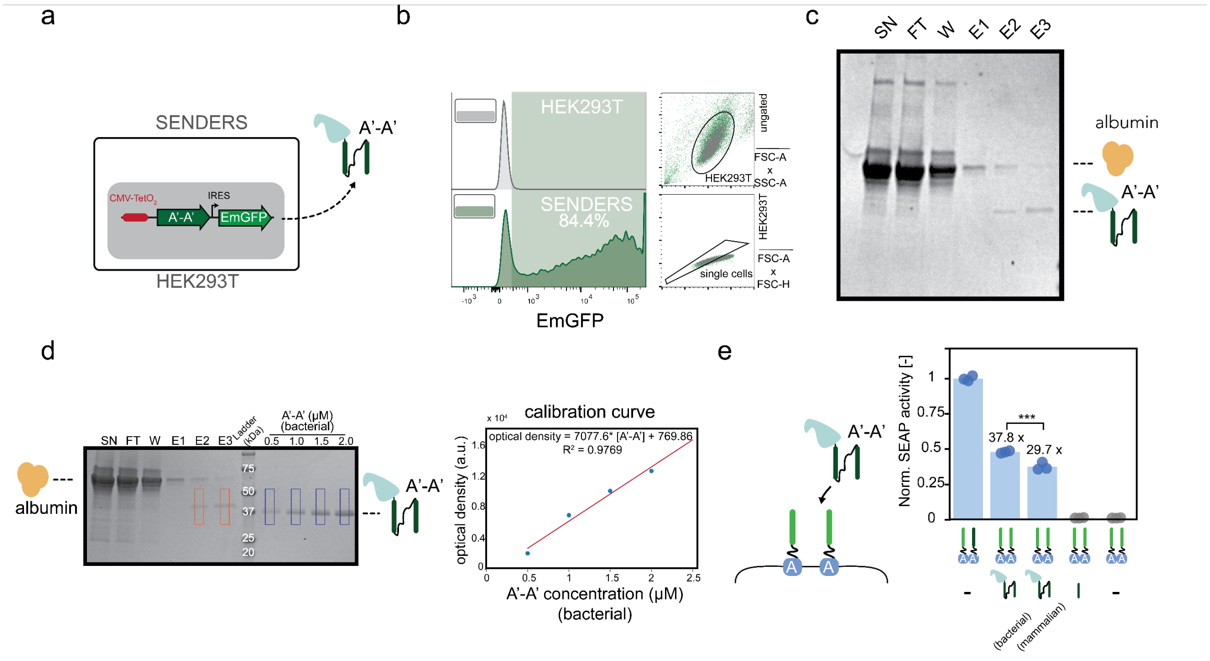


**Supplementary Fig. S17: Engineering of sender HEK293T cells expressing a A'-A' dipeptide. a** HEK293T cells were stably transduced to express and secrete a SUMO-tagged A'-A' bifunctional ligand in the medium (with linker $\mathbf{l}_2$; fused to chicken RPTPσ secretion signal sequence) under the control of CMV-TetO$_2$ promoter. An EmGFP fluorescent protein is also expressed as a bicistron (see Methods and Supplementary Figure S16). **b** Flow cytometry analysis of sender HEK293T cells (green) and control (untransduced HEK293T; grey), showing the gating strategy (dot plots; FSC-A x SSC-A and FSC-H x FSC-H). In the sender population, 84.4% of the cells were EmGFP positive. Backgating is shown on the right. **c** SDS-PAGE analysis, showing the expression of the engineered A'-A' dipeptide expressed in HEK293T mammalian cells. The dipeptide was strep-tag purified from culturing medium that has been in contact with HEK293T cells for four days (see Methods). The presence of albumin is also denoted. **d** Quantification of a A'-A' ligand expressed in mammalian cells, with SDS-PAGE, using gel densitometry analysis (see Methods). SDS-PAGE analysis showing the expressed A'-A' dipeptide (orange rectangles; in E1 and E2) as well as a titration of four known concentrations of A'-A' dipeptide expressed in bacteria (blue rectangles). The presence of albumin is also denoted. A calibration curve was obtained by estimating the optical density (a.u.) of 0.5, 1, 1.5 and 2 μM A'-A' dipeptide ligand expressed in bacteria and fitting the data using linear regression (see Methods). optical density = 7077.6 * [A'-A'] + 769.86, $R^2$=0.9769. **e** Normalized SEAP activity in HEK293T cells transiently transfected with A-type receptor with linker $\boldsymbol{\alpha}_2$ (8 aa, GS repeats), and incubated with 0.12 μM SUMO-tagged purified A'-A' ligand expressed in bacterial or mammalian cells or a monomeric A' CC for 48 hours (see Methods). Bars indicate mean activity; individual data points represent independent triplicates. Normalization is done on the SEAP expression of the A:A' receptor heterodimer with linker length **$\alpha_2$** of eight aa (most left bar). IRES: internal ribosome entry site, SN: supernatant, FT: flow through, W: wash fraction, E1: elution fraction 1, E2: elution fraction 2, E3: elution fraction 3. Significance (ANOVA with Bonferroni test, see Supplementary Table S6) is noted above bars. ns: $p>0.05$, *p $\leq 0.05$, **p $\leq 0.01$, ***p $\leq 0.001$. Fold change compared to cells incubated with A' monomer is noted above the bars.

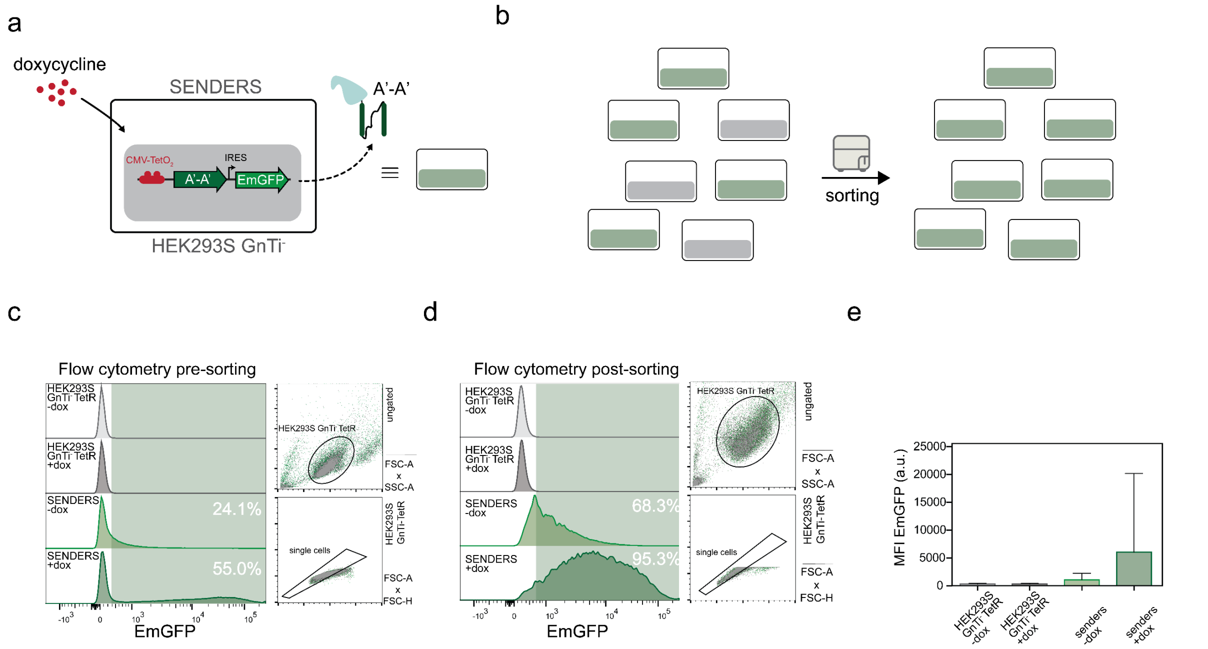


**Supplementary Fig. S18: Engineering of sender HEK293S GnTi⁻ TetR cells expressing a A'-A' dipeptide. a** HEK293S Gnti⁻ TetR cells were stably transduced to, upon induction with doxycycline (dox), secrete a SUMO-tagged A'-A' bifunctional ligand (with linker **$l_2$**) fused to chicken RPTPσ secretion signal sequence, under the control of CMV-TetO₂ promoter. An EmGFP fluorescent protein is also expressed as a bicistron (see Methods and Supplementary Figure S16). **b** To enrich the population of cells expressing the gene, a polyclonal population of cells were sorted based on EmGFP intensity (see Methods). **c** Flow cytometry analysis of sender HEK293S Gnti⁻ TetR cells (green) and control (untransduced HEK293S GnTi⁻ TetR cells; grey) prior to sorting, showing the gating strategy on the right (dot plots; FSC-A x SSC-A and FSC-A x FSC-H). In the sender population and without the addition of dox 24.1% of the cells were EmGFP positive. The addition of dox results in 55.0% EmGFP positive cells. Backgating is shown on the right. **d** Flow cytometry analysis of sender HEK293S Gnti⁻ TetR cells (green) and control (untransduced HEK293S GnTi⁻ TetR cells; grey) following sorting (see Methods). In the sender population and without the addition of dox 68.3% of the cells were EmGFP positive. The addition of dox results in 95.3% EmGFP positive cells. Backgating is shown on the right. **e** Median fluorescent intensity (MFI; a.u.) with robust standard deviation (rSD) of EmGFP of control and sender cells with and without the addition of dox. IRES: internal ribosome entry site.

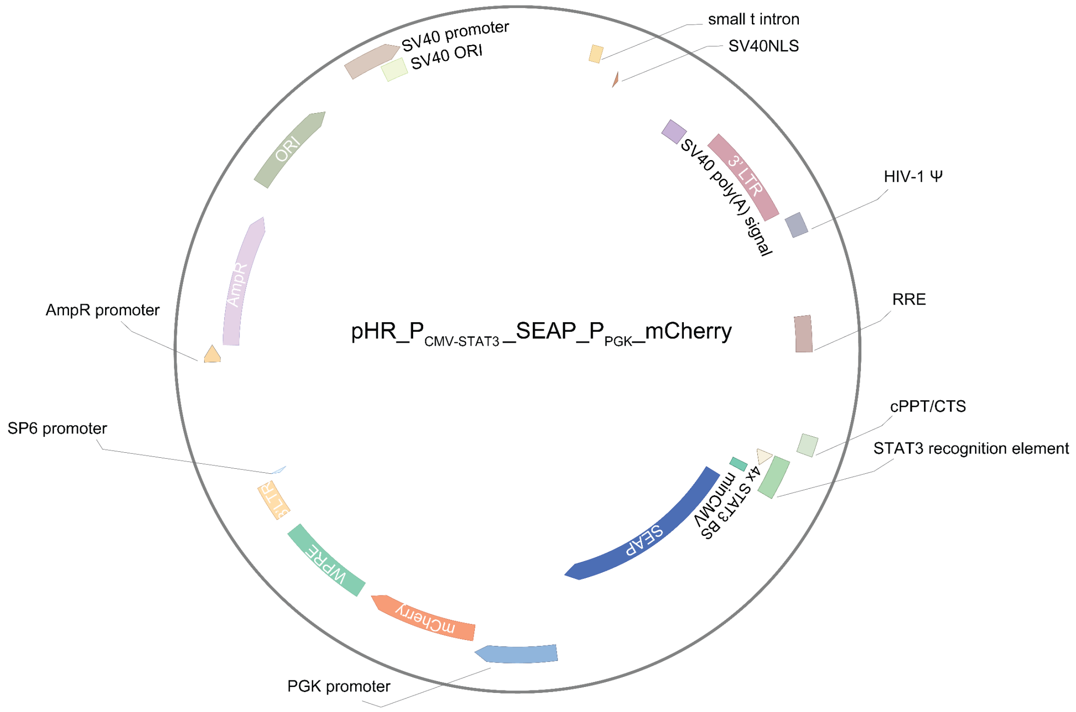


**Supplementary Fig. S19: Plasmid map and sequence of lentiviral expression vector expressing the SEAP reporter gene. a** Plasmid map of pHR vector (see Supplementary Table S3) containing the reporter secreted alkaline phosphatase (SEAP) gene. The reporter is controlled by a signal transducer and activator of transcription 3 (STAT3) recognition element (see Supplementary Table S8), which consists of a minimal cytomegalovirus promoter (minCMV) and four STAT3 binding sites (4x STAT3 BS). A phosphoglycerate kinase (PGK) promoter controls the expression of mCherry fluorescent protein. WPRE: Woodchuck Hepatitis Virus posttranscriptional regulatory element, LTR: long terminal repeat, AmpR: ampicillin resistance, ORI: origin of replication, SV40: Simian vacuolating virus 40, HIV-1 ψ: retroviral psi packaging element, RRE: Rev response element, cPPT/CTS: central polypurine tract/central termination sequence.

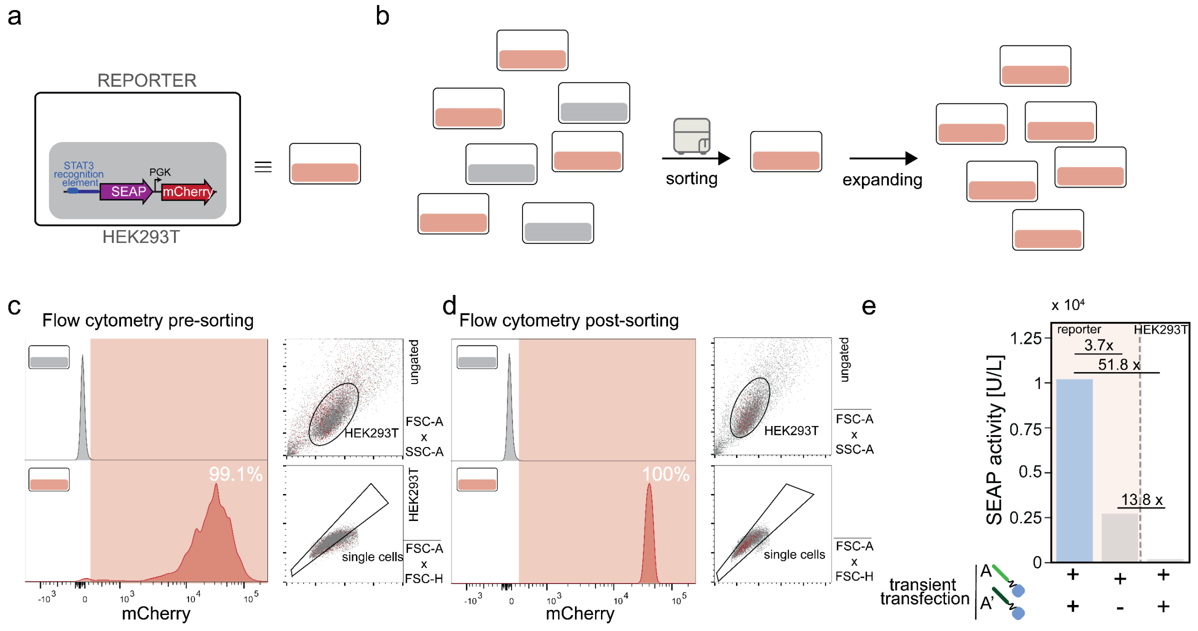

**Supplementary Fig. S20: Engineering of SEAP reporter HEK293T cells. a** HEK293T cells were transduced to express the secreted alkaline phosphatase (SEAP) reporter gene, under the control of a signal transducer and activator of transcription 3 (STAT3) recognition element (see Supplementary Figure S19 and Supplementary Table S8). An mCherry fluorescent protein is expressed under the control of a constitutive phosphoglycerate kinase (PGK) promoter. **b** Monoclonal reporter cell lines were obtained by sorting single cells based on mCherry fluorescence and subsequent expanding (see Methods). **c** Flow cytometry analysis of reporter HEK293T cells (red) and control (untransduced HEK293T cells; grey) prior to sorting, showing the gating strategy on the right (dot plots; FSC-A x SSC-A and FSC-A x FSC-H). Of the reporter population, 99.1% cells were mCherry positive. **d** Flow cytometry analysis of a monoclonal reporter HEK293T cell population (red) and control (untransduced HEK293T cells; grey) after sorting and expanding (see Methods). All reporter cells were positive for mCherry. Backgating is shown on the right. **e** SEAP activity (in U/L) in reporter cells or HEK293T control transiently transfected with A-A' receptor heterodimer and A-type receptors with linker **$\alpha_2$** of 8 aa (GS repeats). Receptor heterodimerization resulted in a 3.7-fold increase of SEAP expression. Bars indicate mean activity (n=1). Fold increase is denoted above bars.

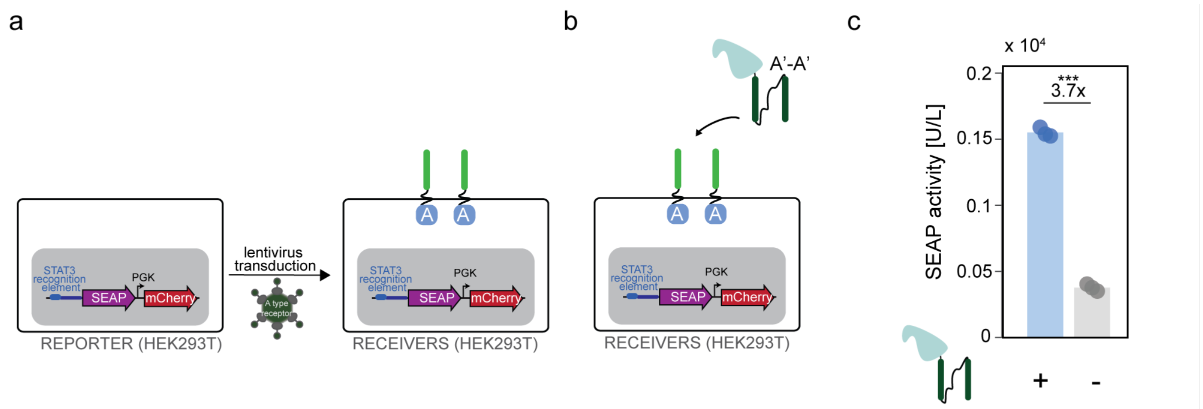


**Supplementary Fig. S21: Engineering HEK293T receiver cells expressing receptor and reporter. a** To engineer receiver cell lines, HEK293T reporter cells (see Supplementary Figure S20 and Methods) were stably transduced to express A type receptor with linker **$\alpha_2$**. **b** Receiver cells were incubated with 0.12 µM cognate SUMO-tagged dipeptide A'-A' ligand (expressed in bacteria; see Methods). **c** SEAP activity (in U/L) in receiver cells incubated with SUMO-tagged dipeptide A'-A' ligand with linker **$l_2$**. Ligand induced receptor dimerization resulted in a 3.7-fold increase of SEAP expression. Bars indicate mean activity; individual data points represent independent triplicates. Significance (t-test) is noted above bars (see Supplementary Table S5). ns: p>0.05, *p ≤0.05, **p ≤0.01, ***p ≤0.001.

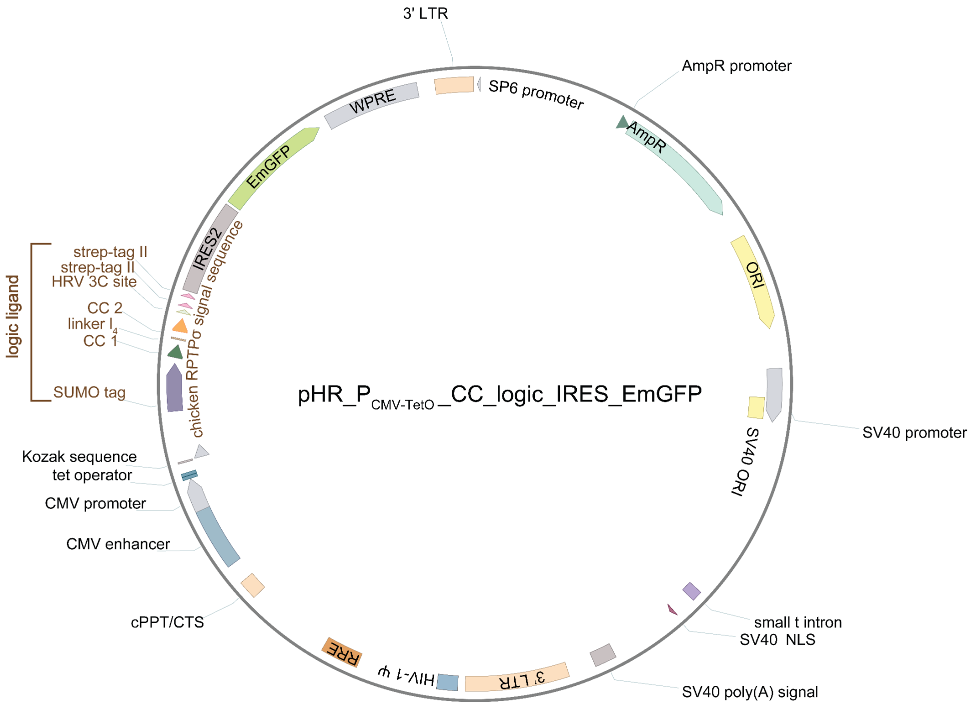


**Supplementary Fig. S22: Plasmid map and sequence of pHR expression vector expressing A'-Γ and Γ'-A' ditopic ligands for AND gate logic. a** Plasmid map of lentiviral vector containing the A'-Γ or Γ'-A' bifunctional ligand (denoted here as logic ligand; in brown). The ligand consists of a chicken RPTPσ signal sequence for secretion, a SUMO-tag, CC 1 (A' CC, see Supplementary Table S1), a linker **l**$_4$ (Supplementary Table S7), CC 2 (Γ or Γ' CC Supplementary Table S1) and a twin strep-tag for subsequent purification (see Methods). The construct is under the control of a mammalian human cytomegalovirus (CMV) enhancer and promoter and two *TetO* operator sequences[3]. The construct is followed by an internal ribosome entry site (IRES) from encephalomyocarditis virus (EMCV) followed by an emerald green fluorescent protein (EmGFP); allowing bicistronic expression of the dipeptide and EmGFP. WPRE: Woodchuck Hepatitis Virus posttranscriptional regulatory element, LTR: long terminal repeat, AmpR: ampicillin resistance, ORI: origin of replication, SV40: Simian vacuolating virus 40, HIV-1 ψ: retroviral psi packaging element, RRE: Rev response element, cPPT/CTS: central polypurine tract/central termination sequence.

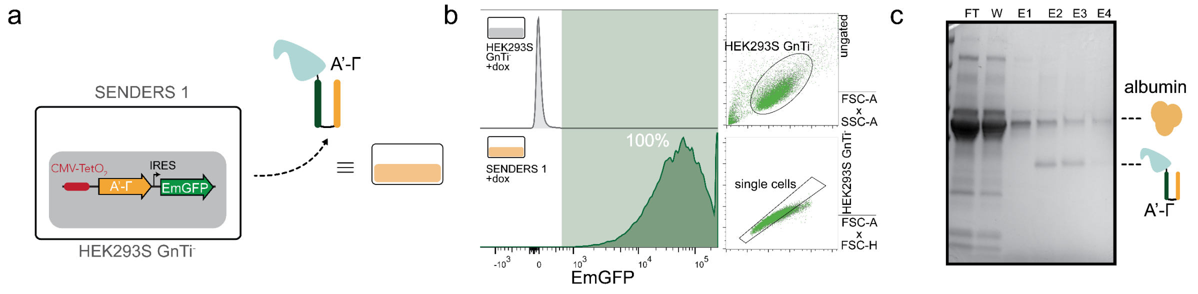


**Supplementary Fig. S23: Engineering of sender HEK293S GnTi$^-$ cells expressing A'-Γ dipeptide. a** HEK293S Gnti$^-$ cells were stably transduced to excrete a SUMO-tagged A'-Γ bifunctional ligand (with linker **l**$_4$; fused to chicken RPTPσ secretion signal sequence) under the control of CMV-TetO$_2$ promoter (senders 1). An EmGFP fluorescent protein is also expressed as a bicistron (see Methods and Supplementary Figure S22). **b** Flow cytometry analysis of sender 1 cells (green) and control (untransduced HEK293S GnTi$^-$ cells; grey) showing the gating strategy on the right (dot plots; FSC-A x SSC-A and FSC-A x FSC-H). **c** SDS-PAGE analysis, showing the expression of the engineered A'-Γ dipeptide expressed in sender 1 cells. The dipeptide was Strep-tag purified from culturing medium, containing dox (see Methods). The presence of albumin is also denoted. IRES: internal ribosome entry site. FT: flow through, W: wash fraction, E1: elution fraction 1, E2: elution fraction 2, E3: elution fraction 3, E4: elution fraction 4.

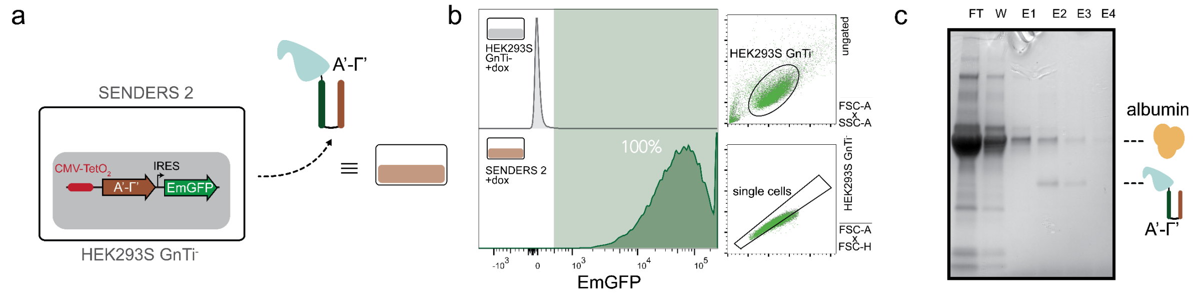


**Supplementary Fig. S24: Engineering of sender HEK293S GnTi⁻ cells expressing a Γ'-A' dipeptide. a** HEK293S Gnti⁻ cells were stably transduced to excrete a SUMO-tagged A'-Γ' bifunctional ligand (with linker **l₄**; fused to chicken RPTPσ secretion signal sequence) under the control of CMV-TetO₂ promoter (senders 2). An EmGFP fluorescent protein is also expressed as a bicistron (see Methods and Supplementary Figure S21). **b** Flow cytometry analysis of sender 2 cells (green) and control (untransduced HEK293S GnTi⁻ cells; grey) showing the gating strategy on the right (dot plots; FSC-A x SSC-A and FSC-A x FSC-H). **c** SDS-PAGE analysis, showing the expression of the engineered A'-Γ' dipeptide expressed in sender 2 cells. The dipeptide was Strep-tag purified from culturing medium, containing dox (see Methods). The presence of albumin is also denoted. **d** Native-PAGE analysis of purified A'-Γ' ligand. IRES: internal ribosome entry site. FT: flow through, W: wash fraction, E1: elution fraction 1, E2: elution fraction 2, E3: elution fraction 3, E4: elution fraction 4.

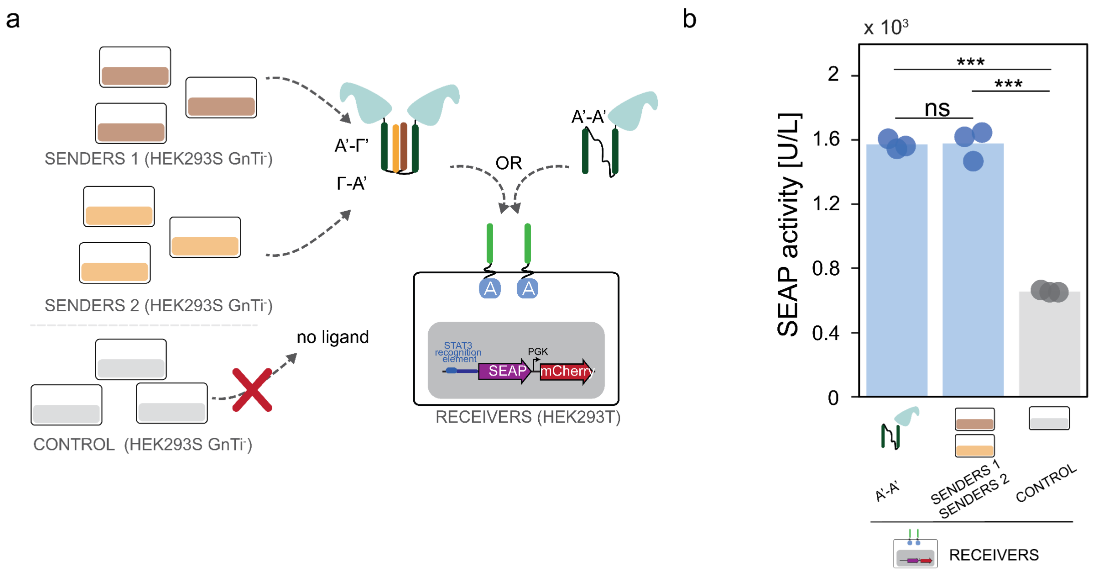


**Supplementary Fig. S25: AND gate logic achieves equal levels of receptor activation as incubation with cognate ligand. a** Receiver cells (Supplementary Figure S21) were incubated with either ditopic A'-A' ligand or were co-cultured with Sender 1 (Supplementary Figure S23) and Sender 2 (Supplementary Figure S24) or control (untransduced HEK293S GnTi⁻ cells). **b** SEAP activity (in U/L) receivers (3.6 x $10^5$ cells) incubated with 0.12 µM A'-A' ligand or co-cultured with senders 1 (1.2 x $10^5$ cells) and senders 2 (1.2 x $10^5$ cells) or control cells (2.4 x $10^5$ cells) for 48 hours (see Methods). Bars indicate mean activity; individual data points represent independent triplicates. Significance (one way ANOVA with Tukey's multiple comparison test) is noted above the bars. ns: $p>0.05$, *$p \leq 0.05$, **$p \leq 0.01$, ***$p \leq 0.001$. (see Supplementary Table S6).

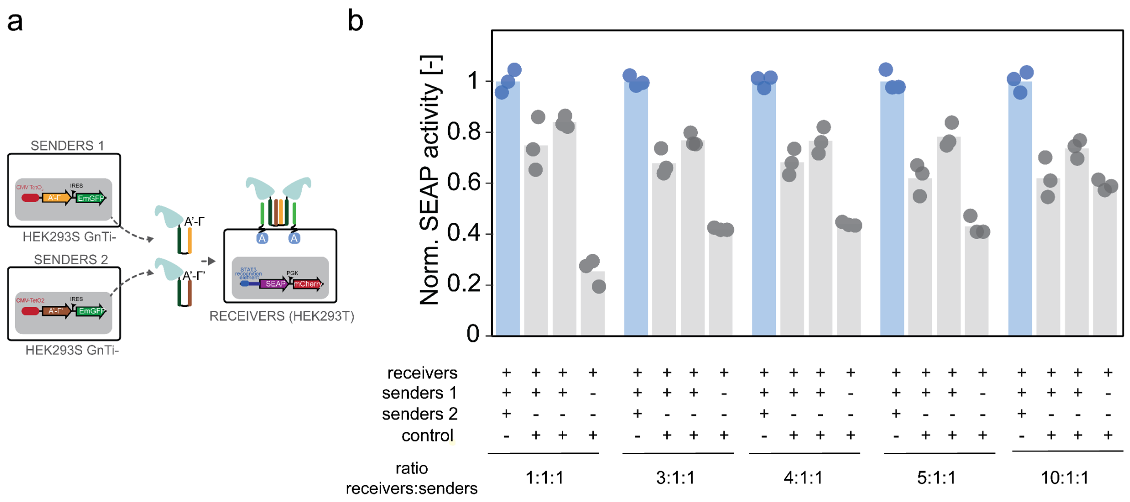


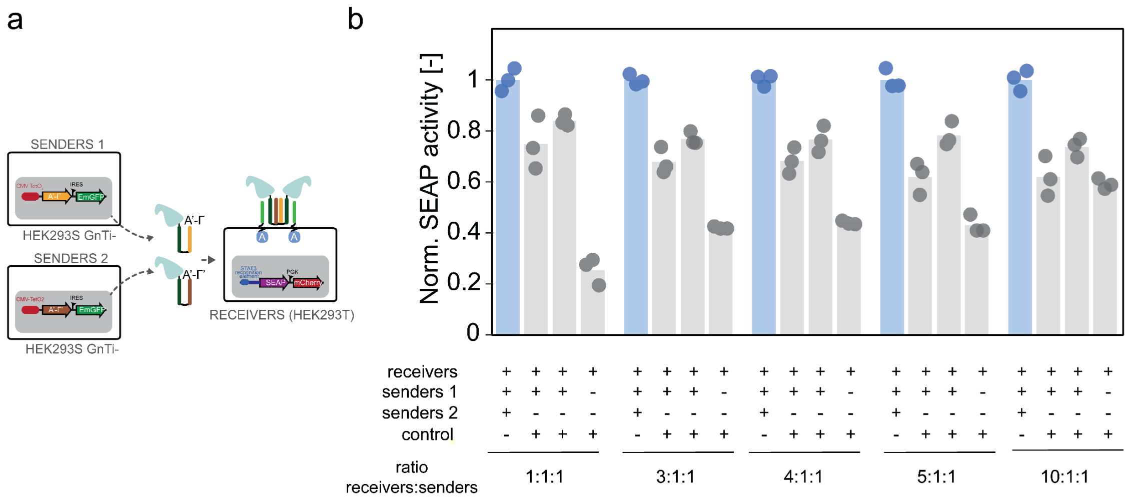


**Supplementary Fig. S26: AND gate logic for different ratios of receiver : sender populations. a** Receiver cells (Supplementary Figure S21) were co-cultured with Sender 1 (Supplementary Figure S23) and Sender 2 (Supplementary Figure S24) or control (untransduced HEK293S GnTi$^{-}$ cells) in a variety of receiver : sender 1 : sender 2 ratios (see Methods and Supplementary Table S10). **b** Normalized SEAP activity in co-culture of receivers and senders 1 and/or senders 2 or control cells (see Methods). Bars indicate mean activity; individual data points represent independent triplicates. For all different conditions, normalization is done on the condition where receiver cells were incubated with both senders 1 and senders 2 (blue bars).
